# Super-resolution visualization of distinct stalled and broken replication fork structures

**DOI:** 10.1101/2020.01.19.912014

**Authors:** Donna R. Whelan, Wei Ting C. Lee, Frances Marks, Yu Tina Kong, Yandong Yin, Eli Rothenberg

**Author notes:** Corresponding Authors Donna R Whelan, PO 199, Pharmacy & Biomedical Sciences La Trobe University, Bendigo, VIC, 3552, Eli Rothenberg, 550 1st Avenue MSB 394 New York, NY 10016.

## Abstract

Endogenous genotoxic stress occurs in healthy cells due to competition between DNA replication machinery, and transcription and topographic relaxation processes. This causes replication fork (RF) stalling and regression, which can further collapse to form single-ended double strand breaks (seDSBs). To avoid mutagenesis, these breaks require repair via Homologous Recombination (HR). Here we apply multicolor single molecule super resolution microscopy to visualize individual RFs under mild stress from the trapping of Topoisomerase I cleavage complexes, a damage induction which closely mimics endogenous replicative stress. We identify RAD51 and RAD52, alongside RECQ1, as the first responder proteins to stalled but unbroken forks, whereas Ku and MRE11 are initially recruited to seDSBs. Ku loads directly onto the DSB end whereas MRE11 associates with nascent DNA away from the break, and both proteins colocalize contemporaneously with a single seDSB. We are thus able to discern closely related RF stress motifs and their repair pathways *in vivo,* uncovering mechanistic insights into the nature of RF damage and repair.

## Introduction

DNA DSBs are highly genotoxic lesions caused by both exogenous agents and endogenous events related to replication and transcription. Repair of these lesions is predominantly carried out via either non-homologous end joining (NHEJ) or homologous recombination (HR), with the latter pathway requiring a homologous sequence as a template for repair [1, 2]. Recently, the existence and conversion of arrested RFs into either regressed ‘chicken foot’ motifs containing a Holliday junction, or single-ended DSBs (seDSBs) (Figure 1A) has been unambiguously demonstrated in higher eukaryotic cells [3–5]. Specifically, it has been shown that upon mild replication stress (for example, clinically induced by Topoisomerase I (TopI) inhibiting anti-cancer drugs), replication forks (RFs) rapidly slow and reverse rather than undergo an inevitable collision with lesions ahead of the fork [6]. This contrasts with another common stressed RF model which involves stalling of the polymerases with continued unwinding resulting in ssDNA generation[7]. This motif can be better mimicked using hydroxyurea [8, 9], which depletes synthesis substrates – deoxyribonucleotides. Conversion of polymerase-stalled RF motifs into regressed forks has not yet been observed and it is understood that conversion of HU-stalled RFs into DSBs requires extended and/or elevated levels of stress [7, 9]. This is in contrast to CPT-stalled RFs, which better mimic endogenous DSB-causing lesions[5]. Which is to say that an individual RF colliding with a single stranded break or physical blockage, can result in a DSB, whereas, depletion of substrates cannot, in and of itself, generate these breaks. For this reason, historically, CPT has been used to generate DSBs whereas mild HU treatment has been used to generate fork stalling, although more recent work has demonstrated the interconnected complexity of the various RF stress motifs [4, 10].

**Figure 1.**
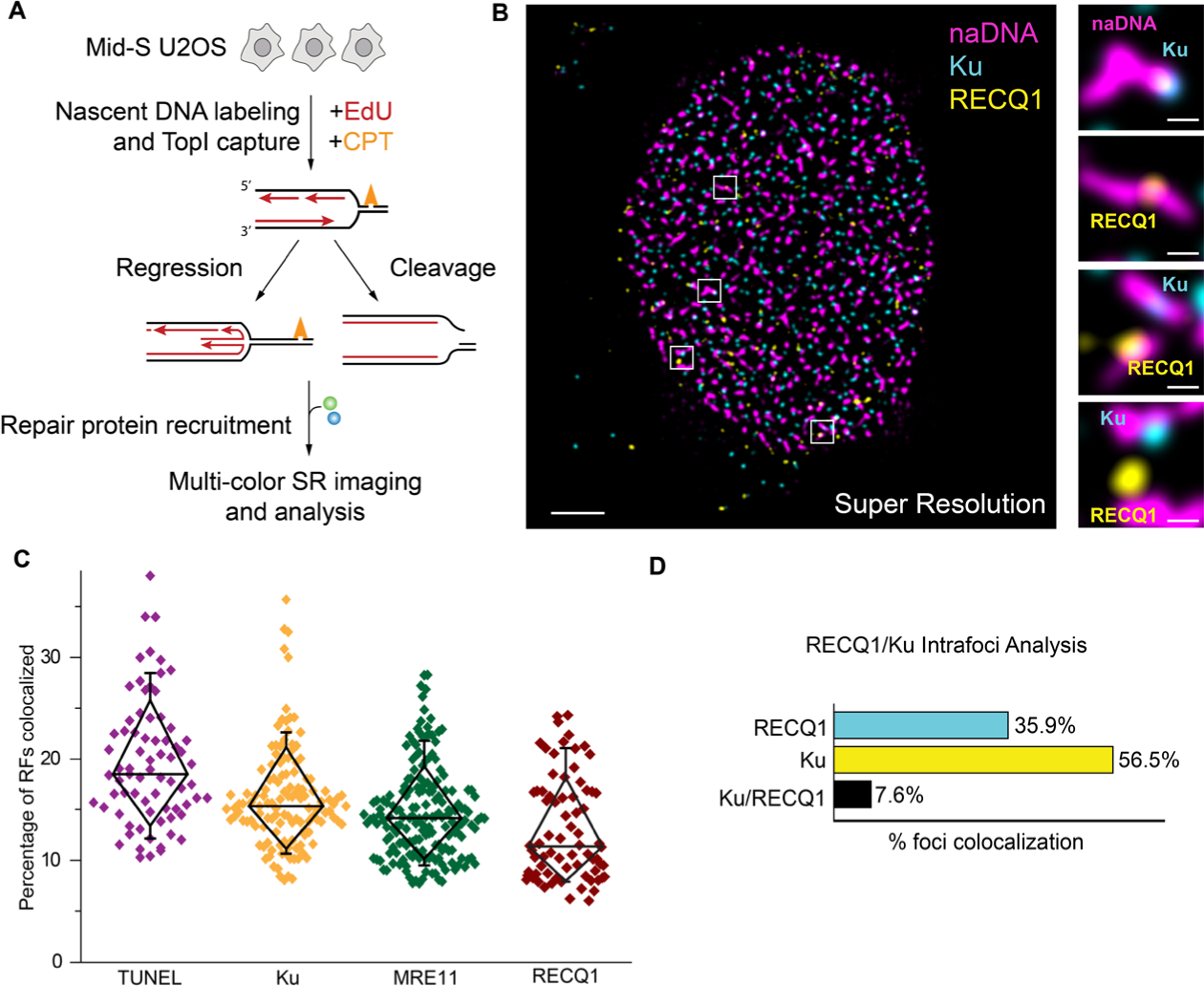
Super-resolution visualization of CPT-induced damaged replication foci. (A) Experimental scheme used to generate and visualize the two hypothesised damage motifs caused by replication stress. (B) Representative super-resolution image of a single nucleus damaged with 100 nM CPT and labelled for naDNA (using EdU, magenta), Ku (cyan) and RECQ1 (yellow). Representative zoomed in images showing either Ku or RECQ1 localized to naDNA foci. Scale bars = 5 μm in whole cell image, 500 nm in zoomed sections. (C) Quantification of the total percentage of RF foci associated with DSBs (via TUNEL signal), Ku, MRE11 and RECQ1 in CPT-damaged cells. (D) Three color analysis showing the exclusionary relationship between RECQ1 and Ku at repair foci. Complete N values available in Table S1. All graphs show mean ± s.e.m. Student’s t-test results shown for comparison with control levels: ns depicts p>0.05, *** depicts p<0.001, **** depicts p<0.0001.

With the recent discovery of RF regression in metazoan cells, it is now hypothesized that fork reversal occurs as a specific means, in the event of upstream lesions, to avoid DSB induction, a type of damage that is fundamentally more complicated to repair than damaged but unbroken RFs. In cases of endogenous DSB induction, even in the absence of stress-inducing drugs, lesions ahead of the RF, such as those caused by processes involving single strand break induction including transcription and topographical relaxation, result in characteristic seDSBs that require specific repair by HR [11–14].

A range of proteins has been identified that regulates and contributes to the HR pathway, and a progression of functional interactions has been proposed to describe it [15–17]. Recently, we used single molecule super resolution (SR) imaging to visualize several of the key steps of this process [18]. Several of the proteins involved in HR are now believed to play protective, reparative, or even antagonistic roles at unbroken but stalled/regressed RFs [19–22]. Following generation of a DSB, either the HR or NHEJ repair pathway must be selected, and there are clear preferences for one or the other depending on cell cycle status, pathway deficiency, and DSB end chemistry [23–25]. However, the mechanisms of DSB repair pathway choice remain poorly understood [26]. This is in part because interactions between, and competition among, the NHEJ protein Ku and initial HR proteins such as MRE11 (as part of the MRN complex), BRCA1, CtIP, and others have not been definitively characterized *in vivo*. MRN, comprising MRE11-RAD50-NBS1, is known to be critical for HR and has been shown to localize immediately to DSB sites with a high affinity for both double stranded DNA and DSBs [27]. This complex is responsible for recruitment of other nucleases, helicases, and mediators, which together orchestrate the extensive resection required at DSBs to generate long 3’ ssDNA overhangs. This resection is an accepted hallmark of canonical HR, and commits the break to HR repair [27, 28]. In contrast, the Ku70/Ku80 ring-motif heterodimer (Ku) is understood to cap the DSB end, blocking exonucleolytic resection and stimulating NHEJ repair [29]. How these two responses to DSBs compete and dictate the repair pathway remains unclear, particularly as both MRN and Ku interactions with DSBs are independent of much of the DNA damage response-signaling cascade and lack clear recruitment pathways [2]. MRE11 has also been identified as both an antagonist and a mediator of stalled RF protection, with some studies demonstrating BRCA2- and PARP1-dependent RF protection from MRE11 degradation [8, 20, 30]. Conversely, others have argued that recruitment of PARP1 by MRE11 is required for reparative resection [31].

A limiting factor in defining the internal organization and repair progression at stalled/regressed and broken RFs *in vivo* has been the spatial resolution of fluorescence microscopy, a limitation imparted by the diffraction of electromagnetic radiation [32]. This has made it exceptionally difficult to distinguish between single and clustered repair sites, as well as the many proteins involved. Indeed, conventional immunofluorescence imaging routinely requires upwards of 100 colocalized fluorophores to generate discernable foci within the nucleus [33], often necessitating the use of clustered or overly potent damage induction. Single molecule localization SR imaging confers a ten-fold improvement in image resolution over conventional fluorescence microscopy [32, 34] together with single-molecule sensitivity, and thus affords the possibility of examining single DSB sites *in vivo* [18]. Here we describe assays based on the *d*STORM variant of SR [35] to achieve this, and their application to determine the specific features and differentiation of early repair responses at single stalled RFs and seDSBs *in vivo*. As a result, we have been able to specifically visualize the initial recognition processes of stressed and broken RFs by first responder proteins, and differentiate between these distinct damage motifs. We found that, along with the repair helicase RECQ1, RAD51 and RAD52 are recruited to stalled but unbroken forks within the first 60 minutes of damage response; this contrasts with the later recruitment of RAD51, RAD52 and BRCA2 to DSBs which we previously found requires 2-4 hours of resection using SR microscopy assays [18]. DSBs recruit both Ku and MRE11 irrespective of their eventual repair pathway choice and do so in a spatially distal organization such that Ku interacts with the DSB end and MRE11 with nearby double-stranded DNA. PARP-inhibition successfully blocks stalled fork protection, converting these damage structures into seDSBs. Our findings represent the first observation and differentiation of individual stalled/regressed versus broken RFs *in vivo* and characterizes the behavior of first responder proteins to these sites.

## Results

### Visualization of distinct replication-associated seDSBs and fork stalling events in individual cells

We used multi-color single molecule SR microscopy [32, 34, 36–38] to determine the spatiotemporal behavior and crosstalk of key ‘first responder’ proteins at individual damaged RFs in human cells (Figure 1A-B). To achieve this, we synchronized U2OS cells in early-mid S-phase and induced a low level of replication stress by treating them with 100 nM of camptothecin (CPT) for one hour [5, 12]. Concurrently, we pulse-labelled nascent DNA (naDNA) using EdU to detect all active RFs [17, 38, 39]. We considered the potential for this simultaneous damage/labelling approach to result in a loss of all EdU incorporation in response to damage to all RFs, or complete cell cycle arrest. Comparison of CPT-treated and control cells, however, did not demonstrate any loss of EdU signal and we further concluded that only a minor proportion of RFs were affected by the low level of CPT damage (Figure 1C). This was in good agreement with previous reports of low DSB induction at similar doses [5, 6]. Furthermore, our observations support recent work that shows that individual RFs exist within the cell spatially distinct from each other, with only pairs of RFs from divergent origin firings forming significant higher order structures during early-mid S phase [40, 41]. Importantly, these findings, in agreement with our own observations, have contradicted the longstanding model of replication organization which postulated the existence of replication factories which contain and coordinate numerous replicons. By costaining pulsed EdU with the replication proteins PCNA and MCM we could well discern individual forks and their directionality (Figure S1). However we do note ongoing contention regarding this issue with new SR studies that describe replication domains in which RFs can be differentiated, but occur in organized structured areas [42]. In light of this ongoing controversy, we again highlight the exceptionally low CPT dose, and its minimal effects on EdU incorporation, as well as protein recruitments (Figure 1C). Finally, we also established that the information obtained in 2D SR projections was representative of, and did not alter conclusions from, flattened 3D SR images (Figure S1).

At low dosages, CPT captures TopI cleavage complexes (TopIcc) ahead of a subpopulation of progressing RFs; this causes them to slow, stall and potentially regress or break to form seDSBs [5]. Therefore, by labeling naDNA, we established a marker for CPT-damaged RFs as we have previously shown [17, 18]. Apart from advantageously positioning DSBs at naDNA foci, CPT is an ideal drug for studying the HR process because the resulting seDSB products closely resemble those generated by endogenous stress from RF-competitive transcription and topographical modification activity [5, 10]. To provide colocalization information, we also developed and validated co-staining assays optimized for SR imaging based on a modified TUNEL assay for enzymatically labelling DSBs [43] as well as immunolabeling protocols for visualizing various key DNA repair proteins.

We used these assays to generate three-color SR images of z-slices of single nuclei in which hundreds of single replication foci could be distinguished and various colocalized proteins quantified. To test our assay, we labeled DSB ends using the TUNEL assay and immunolabelled several key proteins that have been identified as potential first responders in different fork repair pathways, including Ku, MRE11, RAD51, RAD52 and TOPI [44–47]. To specifically detect stalled RFs, we immunolabelled RECQ1 helicase, because it has recently been shown to be involved in rescue of stalled but unbroken RF motifs but has no defined role in DSB repair [45, 46]. Immunostaining of Ku and RECQ1 within the same cells immediately revealed an exclusive relationship in which these two proteins were observed to minimally colocalize with each other at naDNA foci (Figure 1B). Recent literature has demonstrated Ku’s affinity for blunt DSB ends *in vivo* and *in vitro* [29] and hypothesized Ku loading at DSBs *in vivo* irrespective of the eventual repair pathway [47, 48]. In conjunction with our finding that Ku did not colocalize with RECQ1, a known regressed RF repair helicase [46], we identified Ku as a specific marker for seDSBs but not stalled RFs. As a preliminary calculation we quantified the percentage of replication foci colocalized with Ku or RECQ1 or with MRE11, a protein with hypothesized roles at both seDSB [49] and stalled RFs [20, 30], or with DSBs identified using the modified TUNEL assay. This confirmed that only a minority of RFs had been affected by the low dosage of CPT, with between 10 and 20% of replication foci positive for a damage marker (Figure 1C). Furthermore, by only considering naDNA foci that colocalized with either Ku, RECQ1, or both we were able to exclude unaffected RFs (∼75%) and determine that indeed very few damaged naDNA foci colocalized with both Ku and RECQ1 (7.6%) (Figure 1D). This clearly demonstrated the specificity of these two proteins (and their respective antibodies) for the different damage species and showed that naDNA labelling and SR imaging could successfully discern individual stalled RFs from seDSBs *in vivo*, something that has not been achieved previously.

Unexpectedly, the percentage of RFs colocalized with MRE11 was lower than those colocalized with Ku, which appeared to contradict the hypothesis that MRE11 would be detected at both stalled and broken RFs, whereas the TUNEL assay yielded the highest level of occurrence, seemingly indicating that some DSBs had not recruited Ku or MRE11. We considered this an unlikely possibility because without Ku/MRE11 recruitment a DSB would be left unrepaired resulting in gross genomic instability. We reasoned that this observation was more likely due to limitations of the antibody staining, in particular under-staining of individual proteins due to steric hindrance of the antigen, and possible under-sampling because of the super-resolution modality. In contrast, the TUNEL assay appeared highly specific for DSBs within naDNA foci, possibly due to a longer incubation time and the comparatively smaller size (< 3 nm) of the fluorophore conjugate.

To achieve enhanced quantification of the different naDNA proteins association events and to mitigate the potential for over-estimation of colocalization due to the densely populated nuclear environment and cell-to-cell differences in total signal densities, we used a recently established analytical protocol for normalizing the number of detected overlaps to the number of overlaps due to random colocalization [50]. For each single nucleus image, the naDNA and TUNEL/protein foci were defined using automated particle analysis and then the foci in one channel were randomly redistributed throughout the nuclear region of interest (ROI) using a Monte Carlo simulation. The number of overlaps detected in the real nucleus image could then be ratioed to the average number of overlaps generated by the random simulation (taken from 20 simulations) to give a value of 1 if the real-image associations were random and a value above 1 if there was more-than-random colocalization. Based on this analysis, a value of 2 indicated twice as many colocalizations as compared to random, a value of 3 indicated triple, and so on. Importantly, this successfully normalizes cell-to-cell differences in nuclear size as well as the number, size and density of clusters. Furthermore, even if hundreds of colocalizations are present because of the high density of the image, the random simulations will return this same number and the colocalization will be determined insignificant. Such analysis also allows comparison of colocalization in CPT-treated cells with both control levels detected in healthy replicating cells and random levels in simulated data (For details of this and associated analytical protocols for SR data see Figure S2 and [18, 50]).

To test this analysis, we quantified colocalization of TUNEL-labelled DSBs, the phosphorylated histone damage marker γH2AX, and TopI and found that their colocalization increased significantly following CPT damage (Figure 2A-D). The above-random overlap of γH2AX in control cells was expected, particularly as we used U2-OS cells which are prone to increased endogenous replication stress and γH2AX background levels [51]. TopI association with undamaged RFs occurs due to its role relaxing DNA supercoiling ahead of progressing forks [6]. Furthermore, we hypothesized that the detected control level of TUNEL signal might potentially be indicative of non-specificity of the assay at unbroken RFs which called for further experiments and analysis.

**Figure 2.**
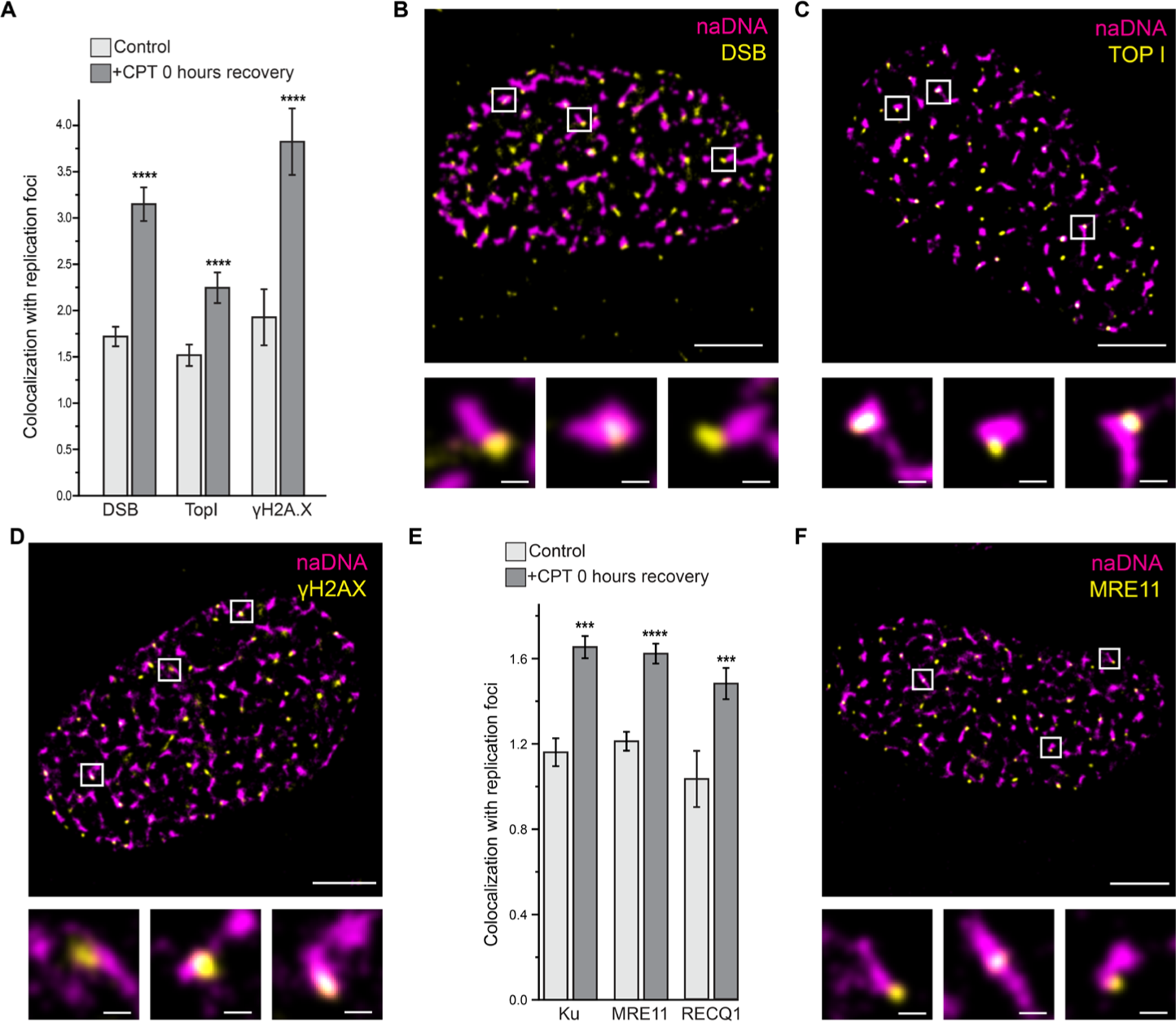
Both fork protection and HR repair proteins colocalize with replication foci following CPT-induced damage. (A) Quantification of colocalization of EdU labelled replication foci (naDNA) with DSBs (via TUNEL labelling), TopI, and γH2AX in undamaged and CPT-damaged cells. (B) Representative SR image of a single nucleus damaged with 100 nM CPT and labelled for naDNA (magenta) and DSBs (using TUNEL)(yellow). Zoomed in regions show colocalization. (C) Representative SR image of a single nucleus damaged with 100 nM CPT and labelled for naDNA (magenta) and TopI (yellow). Zoomed in regions show colocalization. (D) Representative SR image of a single nucleus damaged with 100 nM CPT and labelled for naDNA (magenta) and γH2AX (yellow). Zoomed in regions show colocalization. (E) Quantification of colocalization of replication foci (naDNA) with Ku, MRE11, and RECQ1 in undamaged and CPT-damaged cells. (F) Representative SR image of a single nucleus damaged with 100 nM CPT and labelled for naDNA (magenta) and MRE11 (yellow). Zoomed in regions show colocalization. Complete N values available in Table S1, number of replicates >3, number of cells imaged typically 20-60. All graphs show mean ± s.e.m. Student’s t-test results shown for comparison with control levels: *** depicts p<0.001, **** depicts p<0.0001. Scale bars = 3 μm in whole cell images, 250 nm in zoomed sections.

In contrast to the TUNEL and γH2AX images, the associations of Ku, MRE11 and RECQ1 with naDNA in undamaged control cells were only slightly more than calculated in random simulations, and indeed, equivalent to random localization in the case of RECQ1 (Figure 2E-F). The small amount of MRE11 and Ku colocalization could be explained by the transient binding of these proteins, both of which have affinities for ssDNA, nicks, and gaps that are not associated with DSBs [52, 53]. This result revealed only minimal association between repair proteins and naDNA in undamaged cells and demonstrates the sensitivity of our colocalization analysis. Importantly, quantification of the association of Ku, MRE11 and RECQ1 in CPT-treated cells demonstrated significant recruitment as was expected in response to fork stalling and seDSB induction (Figure 2).

### Ku and MRE11 are contemporaneous and persistent first responders to seDSBs

Ku’s comparable level of recruitment with MRE11 to seDSBs was somewhat surprising given their opposing and competitive roles in DSB repair and a general acceptance that Ku loading at DSBs initiates repair via NHEJ and blocks recruitment of HR proteins [54, 55]. Recent studies have found evidence that on short timescales Ku can be loaded to HR-repaired DSBs [48] and, specifically, to seDSBs, however given the one hour of CPT-damage and evidence that HR nucleases actively remove Ku [47], we expected to detect lower levels of colocalization. Importantly, it is well documented that CPT-treatment generates seDSBs that are repaired by HR because, as well as generating these breaks specifically in S-phase, the single-ended nature of the breaks precludes ligation to an opposing strand. Thus, we concluded that we were observing persistent Ku recruitment to DSBs that would, nonetheless, ultimately be repaired by HR.

In order to focus on the first responder proteins, we confirmed the temporal progression of Ku, MRE11, and RAD51 so as to separate the key canonical steps involved in HR – break recognition, resection, RAD51 nucleoprotein filament formation, homology search and repair. In doing this we aimed to demonstrate that our observations immediately following one hour of CPT-treatment were specific for early HR. We found that although Ku was initially present at seDSBs, within 60 minutes of recovery (and thus potentially 2 hours after break induction) it had been removed (SI Figure 3A). In contrast, MRE11, was recruited to repair foci and remained strongly associated for the first 2 hours of repair during which time naDNA was resected to generate ssDNA (SI Figure 3B). This amount of time for resection, as well as recruitment of various other HR mediators, is in good agreement with previous reports as well as a recent study from our lab [18, 56]. Coinciding with resection, association of RAD51 was detected most strongly at 2-4 hours at which time the RAD51/ssDNA nucleofilament was successfully formed prior to homology search (SI Figure 3C). 12 hours after damage induction, we observed that RAD51 levels had returned almost to control levels, providing an approximate timescale for *in vivo* HR repair, in agreement with our previous observations [18].

**Figure 3.**
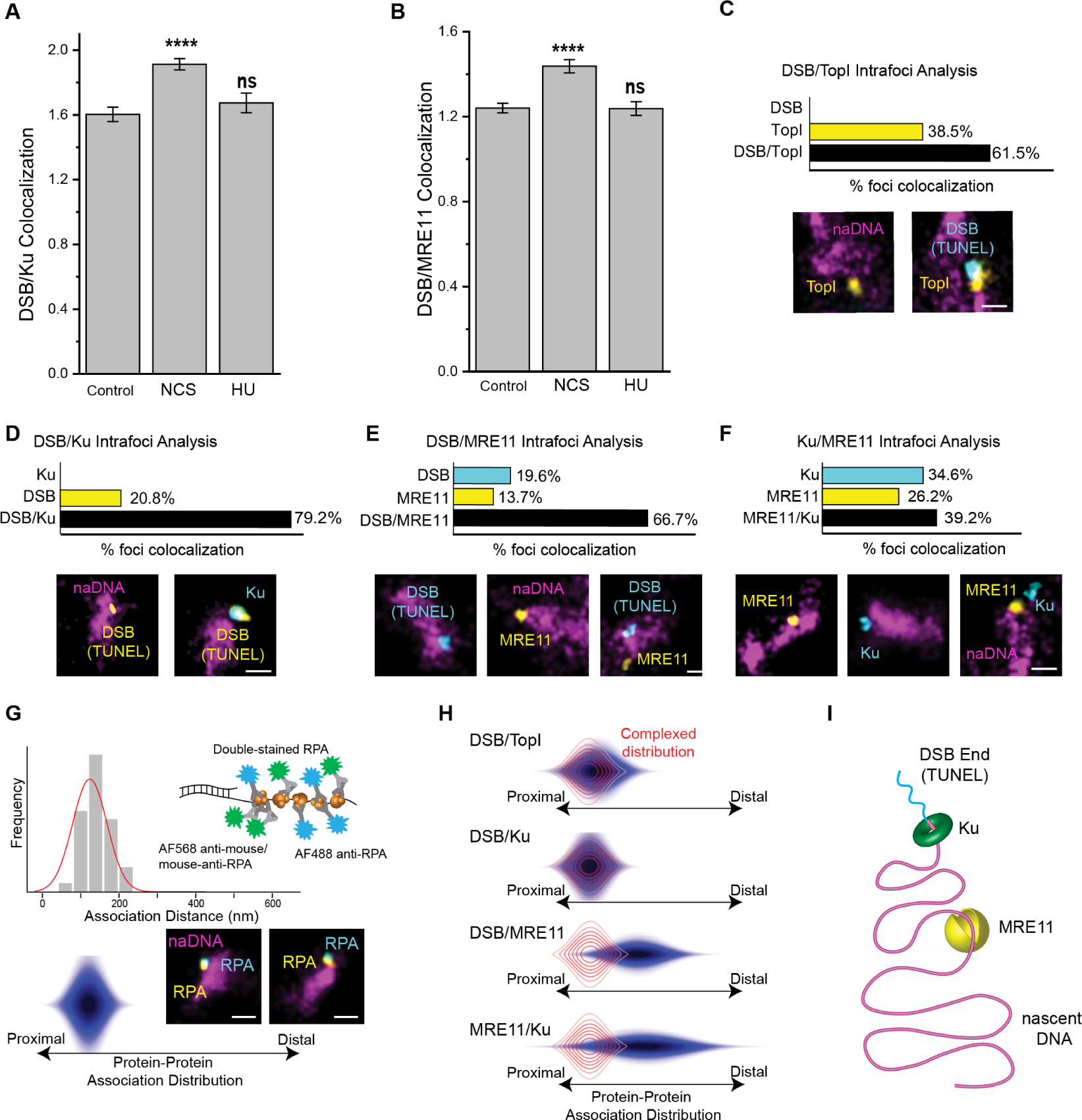
Ku and MRE11 colocalize to seDSB repair foci in a distally separated arrangement with Ku associating with the DSB end and MRE11 with nearby naDNA. (A) Quantification of colocalization of DSBs (via TUNEL labelling) with Ku in control cells and in cells immediately following treatment with NCS or CPT. (B) Quantification of colocalization of DSBs (via TUNEL labelling) with MRE11 in control cells and in cells immediately following treatment with NCS or CPT. (C) Three color analysis of TopI and DSB (using TUNEL labeling) localization at repair foci immediately following 1 hour of CPT damage. (D) Three color analysis of Ku and DSB (TUNEL) localization at repair foci immediately following 1 hour of CPT damage. (E) Three color analysis of MRE11 and DSB (TUNEL) localization at repair foci immediately following 1 hour of CPT damage. (F) Three color analysis of MRE11 and Ku localization at repair foci immediately following 1 hour of CPT damage. (G) A histogram (top) and resulting 3D color-contoured distribution (bottom) and accompanying model (right) showing quantification of the rendered distance between the centers of mass of the same super-resolved RPA foci immunolabeled indirectly with mouse-anti-RPA/Alexa-568-anti-mouse and directly with Alexa-488 conjugated rabbit anti-RPA. These distributions depict proteins closely associated within the foci because of RPA accumulation on ssDNA. (H) Color-contoured protein-protein association distribution maps of protein associations immediately following CPT damage. The red overlap distribution shows the RPA/RPA modelled proximal distribution. (I) A model depicting the association of Ku and MRE11 with a seDSB repair foci in which the TUNEL signal localizes to the break alongside KU while MRE11 localizes distally to double-stranded naDNA. Representative foci shown, scale bars represent 250 nm. Complete N values available in Tables S1-2, number of replicates >3, number of cells imaged typically 20-60, number of individual foci manually scored typically 50-70.

These data confirmed that our observations of Ku and MRE11 localization to RFs immediately following CPT treatment were indicative of initial damage recognition processes. Next we sought to determine the specificity of the TUNEL assay for seDSBs because, although we detected an increase in TUNEL signal following CPT-treatment, we hypothesized that some of this signal could be due to stressed RF species, in particular from stalled and regressed replication forks, because of the high level of colocalization detected in undamaged cells (Figure 2A) [5, 6]. This would present a confounding variable in our use of the TUNEL assay as a seDSB-specific marker for probing the spatial and interdependent relationships of proteins at RF-associated DSBs. Demonstrating that this assay could differentiate between stalled and broken forks would allow further investigation into the key underlying mechanisms of CPT damage – and, indeed, of endogenous DSB-inducing replication stress, namely that the cell employs various complex mechanisms by which to avoid RF collision resulting in DSBs. Previously, it has been shown that this involves slowing, or complete stalling, of the RF, which can traditionally be visualized using pulse/chase DNA fiber analysis, and reversal of the fork to generate a four-way junction, as has more recently been shown using electron microscopy [3, 6, 44, 45]. These stressed RF motifs do not harbor a DSB and so are, presumably, more easily repaired, however, the regressed arm of a RF presents as a blunt end which could, conceivably, be detected by the TUNEL assay and prove difficult to distinguish by SR.

Thus, to establish the specificity of the TUNEL assay and to further verify the lesion-specific detection of our assays, we induced two distinct types of DNA damage, by treating cells with either neocarzinostatin (NCS, 200 ng/ml, 1 hour), which generates two-ended DSBs without inducing replication stress [57], or with a low dose of hydroxyurea (HU, 1 mM, 4 hours), a small molecule drug that causes replication stalling but minimal DSBs under the conditions used [9, 22]. In the case of NCS damage, we were unable to use naDNA as a marker for potential damage sites because NCS-induced DSBs are randomly distributed throughout the nucleus, and are not RF specific. Instead we quantified the colocalization of TUNEL signal with MRE11 and Ku immediately following damage and detected a significant increase demonstrative of NCS-induced DSBs (Figure 3A-B). In contrast, quantification of TUNEL association with both Ku and MRE11 in HU treated cells showed no significant increase which would have indicated off-target TUNEL labelling of stalled RFs and their association with Ku/MRE11 (Figure 3A-B). Furthermore, no colocalization between TUNEL and RECQ1 was detected in NCS-damaged cells (SI Figure 4A). Interestingly a persistent non-random level of TUNEL and TUNEL-associated MRE11 and Ku existed within undamaged cells, possibly due to genuine endogenous levels of genomic stress which are known to be elevated in cancer cells [58, 59]. These observations were further confirmed by quantification of naDNA association with Ku and MRE11 in HU-treated cells, which did not increase with damage because of an absence of DSBs (Figure 3A-B).

**Figure 4.**
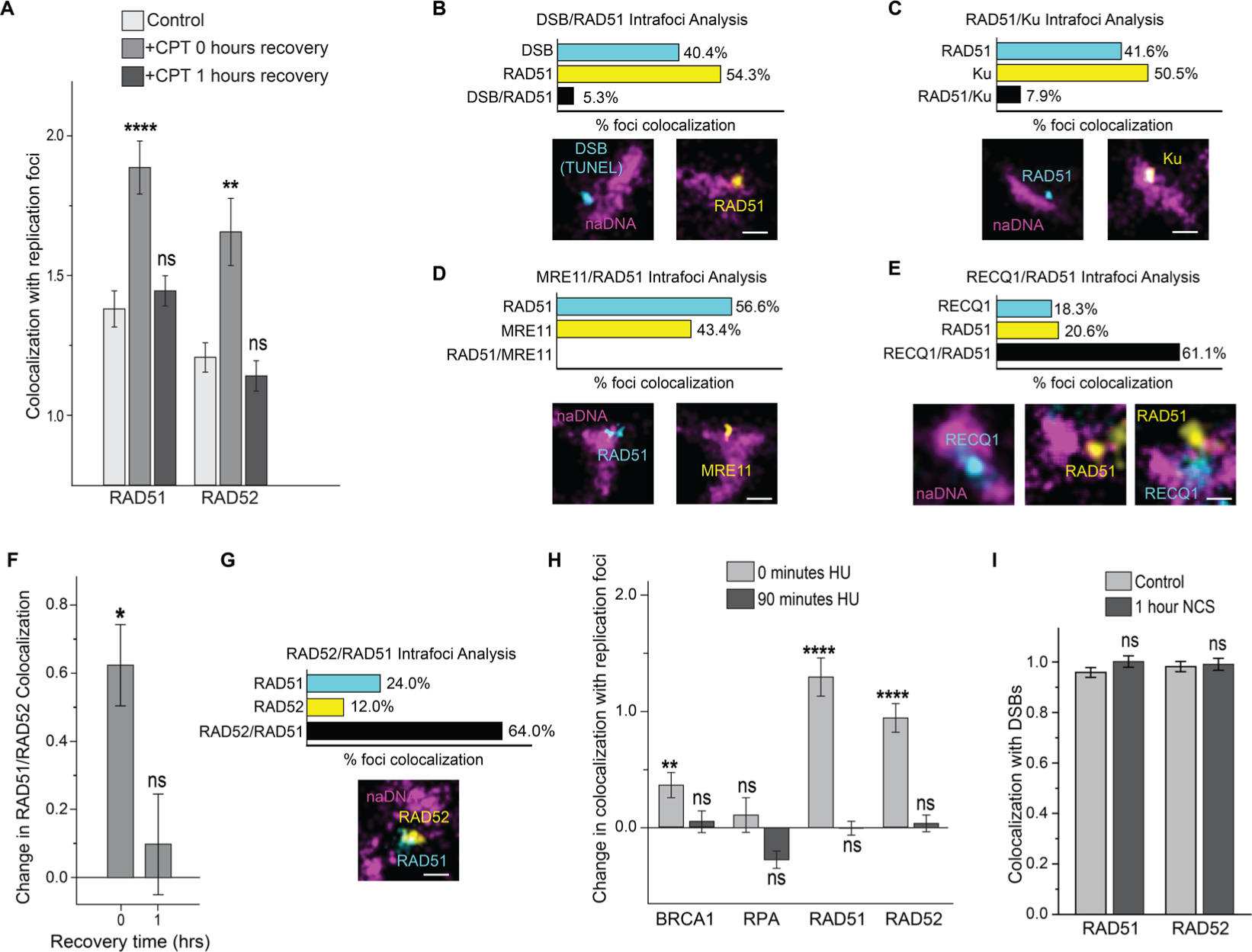
RAD51 and RAD52 are first responders to stalled/regressed RFs. (A) Quantification of RAD51 and RAD52 colocalization with RF foci immediately after CPT damage and following 1 hour of recovery. -C) Analysis of the localization of RAD51 and DSBs, Ku, and MRE11 at RFs following 1 hour of CPT damage. Representative foci shown, scale bars represent 250 nm. (B) Three color analysis of RAD51 and DSB (TUNEL) localization at RF foci immediately following 1 hour of CPT damage. (C) Three color analysis of RAD51 and Ku localization at RF foci immediately following 1 hour of CPT damage. (D) Three color analysis of RAD51 and MRE11 localization at RF foci immediately following 1 hour of CPT damage. (E) Three color analysis of RAD51 and RECQ1 localization at RF foci immediately following 1 hour of CPT damage. (F) Quantification of colocalization of RAD51/RAD52 immediately following 1 hour of CPT treatment and after 1 hour of recovery in normal medium, normalized to control levels. (G) Three color analysis of RAD52 and RAD51 localization at RF foci immediately following 1 hour of CPT damage. (H) Quantification of protein colocalization with naDNA RF foci at unbroken HU-stressed RFs immediately following 4 hours of HU treatment and after 90 minutes of recovery in normal medium from HU treatment, normalized to control levels. (I) Quantification of RAD51 and RAD52 colocalization with DSBs (TUNEL signal) in control and NCS-damaged cells. Representative foci shown, scale bars represent 250 nm. Complete N values available in Tables S1-2, number of replicates >3, number of cells imaged typically 20-60, number of individual foci manually scored typically 50-70. All graphs show mean ± s.e.m. Student’s t-test results shown for comparison with control levels: ns depicts p>0.05, * depicts p<0.05, ** depicts p<0.01, **** depicts p<0.0001.

### Colocalized Ku and MRE11 at seDSBs are distally separated on the DNA

Confident that the TUNEL assay specifically labelled DSBs and not stalled or regressed RFs, and that by fixing cells immediately after one hour of CPT treatment we were visualizing the initial break recognition steps of HR repair, we endeavored to elucidate further spatiotemporal detail. Analysis of three-color images in which both DSBs (via TUNEL) and TopI were labeled alongside naDNA showed that DSB localization to the RF is dependent on the presence of TopI. This result was determined by quantifying the percentage of naDNA foci that were positive for only one or both of the two signals, (SI Figure 2).

Demonstrating the dependence of DSB formation on TopI, we found no colocalization between DSBs with replication foci in the absence of a coinciding TopI signal. In contrast, 38.5% of naDNA-associated TopI was detected without a colocalized DSB, likely because of TopI either trapped or transiently bound ahead of unbroken RFs (Figure 3C). 61.5% of TopI-positive replication foci were also positive for a TUNEL signal, showing a strongly dependent colocalization. Together, these observations demonstrate successful induction of DSBs due to CPT captured TopI complexes colliding with active replication forks.

Next, we considered Ku and MRE11 and their interplay at DSBs. While MRE11 is an integral resection mediator necessary for HR of DSBs, Ku is a core NHEJ protein and its fast unloading and exclusion from HR-fated seDSBs has previously been shown [60, 61]. Nevertheless, we note that the unloading of Ku after one hour is in good agreement with its natural dissociation rate [62] and with recent observations [63] including the blocking of Ku reloading onto minimally resected DNA ends [64].

To investigate the crosstalk of MRE11 and Ku at DSBs, we assessed three-color images to discern their interplay at repair foci. Both MRE11 and Ku were found to colocalize strongly with naDNA associated seDSBs marked by TUNEL, with 66.7% of all Ku positive RFs displaying a DSB signal alongside Ku, while 79.2% of the naDNA foci colocalized with either a DSB or MRE11 demonstrated the coexistence of MRE11 alongside the DSB. Only 20.8% of all DSBs detected lacked a colocalized Ku signal, whereas 19.6% lacked an MRE11 signal. Ku was not detected at naDNA foci in the absence of a break, and only 13.7% of MRE11 association was with RFs lacking a DSB signal (Figure 3D-E). When MRE11 and Ku were co-stained, they were found to associate with repair foci both individually (26.2% and 34.6%, respectively) and together (39.2%), convincingly demonstrating that neither MRE11 nor Ku localization at a single DSB precluded association with the other (Figure 3F).

Because of the high degree of colocalization of Ku with MRE11 at repair foci and their contrary roles in repair pathway choice, we set out to define the internal spatial arrangement of these key proteins within single repair foci by taking advantage of the ten-fold enhanced resolution of our images. To do this, we modelled two-color SR images of closely associated proteins by dual-labelling RPA with Alexa Fluor 568 and 488 in fixed, CPT-damaged cells (Figure 3G) [18]. We then determined the lateral offsets between the two different color signals for more than 50 foci to generate a histogram and 2D protein-protein association distribution map. Because the two different immunolabels of RPA target the same underlying RPA focus, the detected lateral offsets (defined as the protein-protein association distribution) represent protein-protein colocalization that is predominantly proximate and potentially interacting. By similarly determining the distribution maps of pairs of proteins, or DSBs and proteins, colocalized with repair foci, we were able to establish whether they were proximally or distally associated with one another (SI Fig. 2C).

The DSB-TopI association distribution was found to be narrow and spatially proximal, matching closely the distribution determined for dual-stained RPA (Figure 3H, red complexed distribution shown from 3G). This confirmed that DSBs occur close to CPT-trapped TopIccs, and that collision between RFs and single stranded lesions are the predominant cause of the induced DSBs. Similarly, the DSB-Ku spatial distribution demonstrated a close proximal association due to the loading of Ku onto the broken DNA end. In contrast, analysis of the association distribution between both DSBs and MRE11, and MRE11 and Ku, demonstrated distal arrangements in which MRE11 associated with naDNA within the repair foci but away from the seDSB site and Ku (Figure 3H-I). Thus, although MRE11 and Ku routinely colocalize at the same DSB focus, Ku associates directly with the DSB end, while MRE11 localizes distally to naDNA away from the break. This observation is consistent with the endonuclease activity of MRE11, but contradicts current models in which MRE11, as part of the MRN complex, interacts with dsDNA close to the DSB site [23, 48, 49].

### Stalled RFs recruit RAD51 and RAD52 for protection and repair

Although the expected temporal progression of break recognition, resection, and RAD51/ssDNA nucleofilament formation was largely reflected in our data (SI Figure 3), we were surprised to detect small, yet statistically significant, accumulations of RAD51 with replication foci within 60 minutes of CPT treatment (Figure 4A). Even more surprisingly, the observed low levels of RAD51-naDNA association returned to control levels after a further 60 minutes of recovery, but was observed to return to significantly high levels at two hours recovery. This latter time-point is indicative of resection having occurred allowing for construction of the RAD51-ssDNA nueclofilament prior to homology search [18]. While RAD51 is a key protein in HR, nucleofilament formation and recombinase activity have been shown to occur only after extensive resection and RPA coating [28]. Because of this, RAD51 foci formation is usually detected 2-8 hours following DSB induction [65–68]. Moreover, despite a significant increase in colocalization with RF foci immediately following CPT treatment, this could not have represented nucleofilament formation because of its return to control levels after 60 minutes of recovery. In light of recent reports on the interplay of RAD52 with RAD51, we also quantified RAD52 localization to naDNA foci and found that it followed a similar pattern to RAD51 of immediate recruitment before dissipating after one hour. (Figure 4A).

To further elucidate the role of the rapidly recruited RAD51 and RAD52, we analyzed three-color images in which RAD51 was co-stained with naDNA and TUNEL-labeled seDSBs. This revealed that RAD51 was excluded from DSB positive naDNA, with a greater number of RAD51-positive RF foci (54.3%) than DSB-positive foci (40.4%), but minimal colocalization of RAD51 and DSBs (5.3%) (Figure 4B). Similarly, co-staining of RAD51 with either Ku or MRE11 revealed either minimal (7.9%) or no colocalization at replication foci, respectively (Figure 4C-D). We interpret these observations to mean that while Ku and MRE11 colocalize strongly with replication foci harboring seDSBs, the early RAD51 and RAD52 association with replication foci immediately following CPT treatment entails recruitment independent of break formation and canonical HR repair. Strikingly, we found significant colocalization between RAD51 and RECQ1, implicating that the detected early RAD51 recruitment occurs at stalled RFs (Figure 4E).

Despite a lack of well-established role in HR, mammalian RAD52 forms foci during repair and colocalizes with RAD51 foci during late recombinase activity [68, 69]. More recently, we demonstrated that *in vivo* RAD52 acts as the key mediator of early ssDNA/RAD51 interaction, but that it can be replaced by BRCA2 without functional consequence [18]. We therefore considered the possibility of a similar RAD51/RAD52 association and function independent of HR. To test this hypothesis we quantified the change in colocalization between RAD51 and RAD52 immediately following damage and after one hour of recovery, and detected an increase in colocalization at the 0 hour time point but not after 1 hour of recovery, at which time it was found to be at the same level as in control, undamaged cells. This was in agreement with the trends of both proteins’ independent associations with naDNA (Figure 4F). Three-color analysis of these foci confirmed that RAD51/RAD52 colocalized together at naDNA foci immediately following CPT damage (Figure 4G). Based on these observations, we determined that RAD51/RAD52 recruitment to replication foci, while still a consequence of CPT-induced replication stress, occurred specifically at unbroken forks that have stalled or regressed [6, 9, 19, 45].

To further demonstrate this unexpected differentiation in first responder proteins between stalled and broken RFs we again treated cells with HU and assessed colocalization of RAD51, RAD52, RPA and BRCA1 with naDNA [70]. Cell were treated with 1 mM of HU for 4 hours, resulting in replication stress and fork stalling, but minimal fork breakage [9]. Under these conditions we found that RAD51, RAD52 and BRCA1 all localized to RFs (Figure 4H), but upon HU removal, this association diminished within 90 minutes, demonstrating a much faster repair process than canonical HR, which takes several hours. Importantly, we did not detect increased RPA at these foci at either 0 or 90 minutes of recovery following HU treatment, showing the lack of extensive resection required for repair of stalled but unbroken RFs. Thus, notwithstanding questions concerning the role of RAD52 [71–74], these observations supported our hypothesis that both RAD51 and RAD52 are recruited to stalled/regressed RFs.

To contrast HU treated cells, we also assessed cells treated with NCS to generate DSBs in the absence of RF stress. In these cells we found no increased association between DSBs and either RAD51 or RAD52 immediately following one hour of NCS damage (Figure 4I) further demonstrating the early RAD51/RAD52 recruitment to DSBs is uniquely associated with RF stalled/regression. Finally, we assessed early MRE11 association with RAD51 across all drug treatments used here and found no evidence of colocalization, once again revealing their mutually exclusive roles at seDSBs and stalled RFs immediately after low levels of damage induction (SI Figure 4B).

### CPT treatment combined with PARP-inhibition synergistically converts stressed RFs into seDSBs

It has previously been reported that RF protection by stalling/regression is PARP-dependent [6] and more recently, it was discovered that PARP inhibition causes DNA lesions to go unrecognized, blocking the consequent RF stalling/regression that enables repair, instead causing an increase in seDSBs induction [75]. In light of these findings we reasoned that inhibiting PARP in cells that are treated with low dosage CPT would cause the resulting RAD51/RAD52-positive stalled RFs to be bypassed to generate more TopI-induced seDSBs.

To test this hypothesis, we pre-treated cells with the PARP1/2 inhibitor Veliparib for 24 hours before addition of CPT. We used comet assays to determine the level of DSB induction to assess the degree of RF stalling/regression bypass resulting in increased seDSBs in the Veliparib+CPT-treated cells. While both these drugs generated increased levels of DSB induction individually, a synergistic effect was detected upon combination, showing that Veliparib was indeed causing CPT-induced replication stress to result in seDSBs (Figure 5A). This further confirmed the existence of a substantial number of stalled but unbroken RFs in CPT-treated cells. We also assessed Veliparib+CPT-induced DSB induction using SR and found increased levels of DSBs (via TUNEL), MRE11, and Ku association with naDNA foci (Figure 5B).

**Figure 5.**
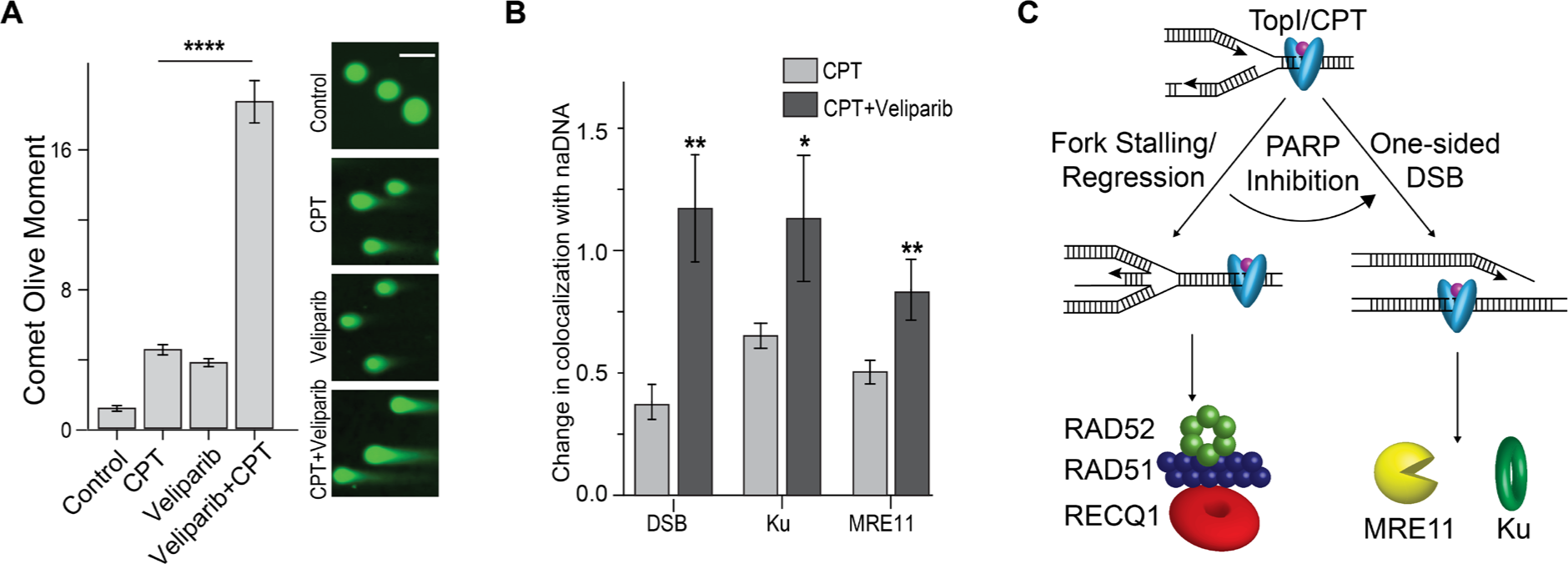
PARP inhibition converts RAD52/RAD51 protected RFs into seDSBs. (A) Comet assays of cells damaged with CPT, Veliparib or both demonstrating synergistic DSB induction upon TopI-induced RF stress combined with PARP inhibition. Representative cells shown. (B) Quantification of DSB (TUNEL), Ku and MRE11 association with naDNA upon PARP inhibition using a combined Veliparib/CPT treatment. (C) A model of the two predominant stress pathways induced by TopIcc: RF stalling/regression, which is protected and repaired quickly and colocalizes with RAD51 and RAD52, and osDSBs, which colocalize with Ku and MRE11 and are repaired via HR. Representative comet assayed cells shown, scale bars represent 15 μm. Complete N values available in Tables S1-2, number of replicates >3, number of cells imaged typically 20-60, number of individual foci manually scored typically 50-70. All graphs show mean ± s.e.m. Student’s t-test results shown for comparison with control levels in (A) and with CPT-only (B): ns depicts p>0.05, * depicts p<0.05, ** depicts p<0.01, **** depicts p<0.0001.

## Discussion

In this study we described powerful single-molecule localization-based assays for differentiating and analyzing distinct RF repair pathways. Using the enhanced spatial resolution and single molecule sensitivity achieved in these assays, we have been able to examine the organization of repair proteins within repair foci consisting of individual seDSBs or stalled/regressed RFs at a comparatively low level of damage induction. This enabled identification of specific, mutually exclusive first responder proteins: RAD51, RAD52, and RECQ1 at stalled forks, and MRE11 and Ku at seDSBs (Figure 5C). Together, these data constitute a direct visualization of closely linked damage motifs *in vivo* revealing hitherto unknown aspects of the specific roles of response proteins. Our observations also contrast findings based on higher damage inductions and biochemical methods which rely on ensemble averaging of cellular data.

The two canonical DSB repair pathways, NHEJ and HR, are both active in the repair of chromosomal breaks during S phase [76]. The cellular abundance of Ku in human cells and its remarkably high affinity for a variety of DSB lesions results in its immediate recruitment to break sites [48]. Despite this, existing models of DSB repair in cells depict two exclusive pathways, each of which proceeds via a set of sequential steps. The data presented here demonstrate the existence of a more refined repair process in which Ku initially loads onto seDSBs generated at collapsed RFs despite their eventual repair via HR. Moreover, we show that MRE11 loads contemporaneously with Ku association, but in a distal position (Figure 3). This contradicts models in which MRE11 initially associates much closer to the DSB and progresses towards the break, suggesting not only that resection is initiated away from the DSB site but that MRE11 remains distal to the break and does not seem to simply execute 5’ to 3’ nucleolytic degradation as previously thought [16, 27]. Instead our findings elucidate the *in vivo* reality of recent *in vitro* studies that showed MRE11 conducting long-range one-dimensional diffusion along dsDNA associated with a DSB [77]. These observations highlight the fact that all cellular DSBs are first bound by Ku, an event that is then followed by repair either via NHEJ or removal of Ku and progression of HR.

A noteworthy strength of our analysis rests on our ability to distinguish specific DDR variability associated with different fork repair processes [17, 78, 79]. This led us to the observation that a subset of RFs colocalized with the HR proteins RAD51 and RAD52 immediately following CPT damage, but did not colocalize with TUNEL signal, Ku or MRE11 (Figure 4). These data are consistent with the notion that these forks do not form seDSBs. This distinctive DDR signature was also observed at forks subjected to replication stress via HU treatment, which specifically causes RF stalling and regression, whereas Ku and MRE11 were not detected in HU-treated cells (Figure 3A-B). Accordingly, we conclude that while low grade CPT treatment results in seDSB formation, it also yields unbroken but arrested RFs that require RAD51/RAD52 to mediate fork protection and regression processes [6]. This conclusion was further strengthened by detection of RECQ1, a helicase known to process stalled RFs (Figure 2E). Importantly, we showed that the activity of RAD51/RAD52 fork protection is PARP1 dependent, with combined Veliparib+CPT treatment resulting in conversion of regressed forks into seDSBs (Figure 5) [6]. This result further shows that endogenous stress caused by transcriptional activity and topographic remodeling ahead of RFs results in both stalled/regressed and broken RFs, and that these are repaired by distinct pathways, each making use of canonical HR proteins.

The novel combination of discriminatory labeling and analytical approaches with state-of-the-art super resolution imaging detailed here has enabled visualization and characterization of key endogenous damage species and subsequent cellular responses *in vivo.* We unequivocally demonstrate the coexistence of both arrested and broken RFs in cells under mild replication stress conditions. We further show recruitment of RECQ1, RAD51 and RAD52 to arrested RFs allowing for expeditious repair, and the loading of Ku and MRE11 specifically onto seDSBs, which are more slowly repaired via HR. The temporal and structural insights derived highlight the capabilities of the assays we describe and identify RAD52 and Ku as having previously unknown roles at stalled RFs and seDSB damage sites, respectively. Our analysis also shows that MRE11 loads and interacts with seDSB DNA away from the break, contrary to current models. Taken together, our observations offer new insights into the roles of various DNA damage species and the response pathways associated with the pathogenesis of cancer, autoimmune diseases, and a range of other genetic disorders.

## Material & Methods

### Cell Synchronization and Drug Treatment

Female human bone osteosarcoma U-2 OS cells (ATCC HTB-96) were routinely cultured in McCoy’s 5A (Modified) medium (ThermoFisher 16600) with 10% FBS (Gemini Bio. 100-106) and 100 U/mL Penicillin-Streptomycin (ThermoFisher 15140) at 37°C and 5% CO_2_.

Cells were seeded on No 1.5 glass coverslips and allowed to adhere in complete medium for 18-24 hours before being switched to FBS free medium for a further 48-72 hours in order to synchronize cells in G0/G1 phase [80]. Cells were subsequently released in complete medium for a further 16 hours to produce an early-mid S phase cell population. To induce RF stress and seDSB generation cells were treated with 100 nM CPT (Abcam 120115) [5] alongside pulse labeling with 10 μM EdU, a thymidine analogue, for one hour so that it would be incorporated into naDNA during replication (Click-iT kit, ThermoFisher C10340) [18]. Cells were fixed immediately following damage or released back into drug-free complete medium for 1-12 hours before later fixation. This allowed for spatiotemporal mapping of slower HR processesControl cells were treated with 0.1% DMSO in place of CPT.

To further perturb and probe the HR process of DSBs, U2OS cells were treated with 100 μM of the PARP 1/2 inhibitor Veliparib (Santa Cruz, 202901) for 24 hours prior to recovery from CPT treatment [81]. To do this, cells seeded on coverslips were administered Veliparib during the final seven hours of serum starvation as well as during release in complete medium and in the CPT/EdU medium. Hydroxyurea experiments were conducted for 1-4 hours at a final concentration of 1 mM, whereas Neocarzinostatin was applied for 1 hour at 200 ng/ml.

### Extraction and Fixation

For clear visualization of chromatin and chromatin-bound nuclear fraction of cells, optimization of fixation and immunolabeling protocols was of paramount importance[82]. In particular, the methods presented here were specifically optimized to achieve a high degree of soluble component extraction while maintaining the ultrastructure of the nucleus and are in good agreement with similar protocols[48]. Pre-extraction of the cells was achieved using room temperature CSK buffer (10 mM Hepes, 300 mM Sucrose, 100 mM NaCl, 3 mM MgCl_2_, and 0.5% Triton X-100, pH = 7.4) for 2-3 minutes with gentle agitation. This was a key step as it removed the majority of the unbound fraction of HR proteins, which decreased the density of proteins within the nucleus and increased the proportion of detected proteins that were directly involved in HR (ie. associated with naDNA at DSB sites). This step also removed much of the cytosolic component, which minimized non-specific binding sites and sources of background and auto-fluorescence. After extraction, cells were fixed in paraformaldehyde (3.7% from 32% EM grade, Electron Microscopy Sciences, 15714) and glutaraldehyde (0.3% from 70% EM grade, Sigma-Aldrich, G7776) in PBS for 15 minutes. The cells were then washed three times with PBS and, if required, stored overnight at 4°C. For fluorescent tagging of the pulse labeled nascent DNA, the copper catalyzed ‘Click’ reaction was used as described in the Click-iT (ThermoFisher, C10640) protocol in conjunction with Alexa Fluor 647 [18]. The cells were blocked with blocking buffer (2% glycine, 2% BSA, 0.2% gelatin, and 50 mM NH_4_Cl in PBS) for 1 hour at room temperature (RT) or overnight at 4°C prior to further staining.

### TUNEL and immunofluorescence labeling

To visualize DSBs, the Click-iT TUNEL kit (ThermoFisher, C10247) was combined with ChromaTide Alexa Fluor 568-5-dUTP (ThermoFisher, C11399) and a modified protocol standardized against established ‘Click’ and Digoxigenin (digoxigenin-11-dUTP: Sigma, 11093088910, anti-digoxigenin: Novus, 31191) protocols [37]. The optimized protocol made use of conjugated AF568-5-dUTP (ThermoFisher, C11399) to label DSBs in conjunction with AF647 ‘Click’ labeled naDNA demonstrating high specificity of TUNEL labels as shown by high levels of overlap with naDNA foci in damaged cells. To achieve this, AF568-5-dUTP was diluted to 50 μM in the EdUTP mix and allowed to react as directed for 12 hours at 37°C. Samples were washed three times in PBS and blocked for a further 30 minutes with blocking buffer.

Antibody labeling of all other proteins was achieved via a combination of direct and indirect labeling with Alexa Fluor 488 and 568 fluorophore labeled antibodies that have previously been validated by as and others for IF experiments. For a complete list, including validating references see SI Table 3.

### Super Resolution Imaging

For SR imaging, coverslips were stored for up to one week at 4°C before being mounted onto a microscope microfluidics chamber just prior to imaging. SR imaging buffer comprising an oxygen scavenging system (1 mg/mL glucose oxidase (SigmaAldrich, G2133), 0.02 mg/mL catalase (SigmaAldrich, C3155), and 10% glucose (SigmaAldrich, G8270)) and 100 mM mercaptoethylamine (Fisher Scientific, BP2664100) in PBS was then prepared fresh and added to the imaging chamber.

A custom-built SR microscope based on a Leica DMI 3000 inverted microscope was used to acquire all the raw data as has been described previously[18]. Briefly, 473 nm (Opto Engine LLC, MBL-473-300 mW), 532 nm (OEM Laser Systems, MLL-III 200 mW) or 556 nm (UltraLasers, MGL-FN-556 200 mW) and 640 nm (OEM Laser Systems, MLL-III 150 mW) laser lines were combined using appropriate dichroics and focused onto the back aperture of a HCX PL APO 100X NA = 1.47 TIRF (Leica) objective via a multi-band dichroic (Chroma, zt405/488/532/640/730rpc, UF1C165837). The incident excitation beam could be translated laterally across the back of the objective to achieve a Highly Inclined and Laminated Optical (HILO) illumination configuration producing better signal-to-noise images. Fluorescence emission was collected through the same objective and dichroic and imaged on either an electron multiplying charge coupled device (EM-CCD) camera (Andor iXon+ 897) or scientific complementary metal-oxide semiconductor (sCMOS) camera (Photometrics, Prime 95B). Fluorescence signal from AF568 and AF647 could be collected simultaneously using a dual-band bandpass filter (Chroma, CY3/CY5, 59007m), and split into two channels on the EMCCD using a dichroic mirror (Semrock, FF660-Di02) in conjugation within a dual-view cube (Photometrics, DV2). AF488 signal was collected subsequently using a narrow single-band filter (Semrock, FF01-531/40). A 405 nm laser line (Applied Scientific Pro., SL-405 nm-300 mW) was introduced to enhance recovery of dark state fluorophores when required. Alternatively, channels were imaged sequentially. S phase nuclei were identified by positive AF647 naDNA signal and 2000 frames at 33 Hz were acquired for each color. Accumulation of more frames increased total signal only marginally (10-20%) and did not affect the overall analysis because immuno-, TUNEL, and naDNA labelling inherently involves multiple fluorophores targeted to the same underlying feature, and because complex structural determinations were not useful to the study. Representative images showing the blinking density and photobleaching can be seen in SI Figure S5. Fitting uncertainty was determined based on photon number, PSF approximation, and the camera noise to be <10 nm in the red and blue channels, and <20 nm in the green channel. Mapping errors based on transformation of bead images and were found to be ∼10 nm for both mapping of the blue channel to the red channel and the green channel to the red channel.

A polynomial morph-type mapping algorithm achieved offset and chromatic correctional mapping of the three channels. To generate the correctional map, each day, prior to imaging, diffraction-limited images of spatially separated broad-range emitting fluorescent beads were acquired across all three colors (Tetraspecks, 100 nm, Life Technologies, T7279). The precise localizations of the beads were obtained by independently fitting their diffraction-limited Point Spread Functions (PSF) with Gaussian functions. The three-color sub-diffraction localizations could then be matched and an elastic mapping matrix generated based on the polynomial morph-type mapping function using an IDL (Exelis Visual Information Solutions) custom mapping script. Three-color raw image stacks of cellular images were then corrected before SR analysis.

### Super Resolution Image Rendering and Analysis

To generate a table of molecular localizations and render SR images for colocalization analysis, the three acquired colors were processed independently using the ImageJ [83] plugin QuickPALM [84] with point spread function fitting constrained to spots with full-width half-maximum of 4-5 pixels (∼264-535 nm) and signal-to-noise ratios better than 3. Images were rendered using 20 nm pixels and recombined to generate three-color micrographs over which masks were manually drawn identifying the nucleus based on the naDNA signal.

To determine the degree of colcoalization, Monte Carlo simulations depicting random distribution of the clusters relative to each other were used as a normalization approach. To achieve this, a random simulation was generated for each experimental image by creating grayscale images of two colors at a time from the image of a nucleus (ie. color-1 and color-2). The two greyscale images were then segmented by automatically applying an Otsu thresholding algorithm to the gray-scale pixels that belonged to the masked nuclear ROI. After cluster segmentation, a random redistribution of one of the colors was simulated by redrawing all of the clusters that belonged to the color-2 experimental image over the static color-1 image. From this ‘randomized’ simulation, both the number and the area of overlap between the two colors were determined.

Using this approach, 20 simulations of random rearrangements were generated for each nucleus in a pairwise fashion, examining Red/Green, Red/Blue, and Green/Blue overlap. The total number or area of overlaps detected in each nucleus (typically 15-100) could then be normalized to the determined random level of overlap expected by dividing the number or area of real overlaps by the average number or area of overlaps in the same simulated nucleus. An average normalized colocalization factor was thus calculated for each pairwise interaction between proteins, seDSB, ssDNA, and naDNA (N typically 30-50 total cells, each comprising 100s of foci, all N values available in Table S1). A colocalization factor of 1 indicated completely random colocalization, while higher factors indicated association and interaction. Colocalization factors were also calculated for undamaged cells to establish control levels. Number of overlaps was used for TUNEL, TopI, Ku, MRE11, and BRCA1 whereas area of overlap was used for γH2AX, RPA, RAD51 and RAD52 due to the expected accumulation of these proteins over multiple time points. In general the trends observed in number of overlaps were comparable to those observed in area but were more striking in the latter case for accumulating proteins. Change in colocalization was further normalized against the degree of overlap detected in undamaged control cells fixed immediately following EdU pulse labeling. For this reason several overlaps show a negative change due to a decrease in protein association when the averaged damage foci is compared to the averaged replicating fork foci. For further explanation see [18, 50].

To assess the prevalence of colocalization, dependence or exclusionary relationships between proteins at DSBs, all foci within a nucleus containing both naDNA and at least one of the two proteins stained were identified and quantified to determine the percentage of foci with colocalized proteins, as opposed to those positive only for one or the other. The average proportions were calculated and depicted in cumulative bar graphs.

To assess the intrafoci organization of proteins at DSBs, the distance between centers-of-mass of colocalized proteins at naDNA was determined. These distances were used to generate a histogram, which could be approximated with a single or double Gaussian. To standardize intrafoci two-color distances descriptive of closely associated proteins, RPA was double labeled with AF568 and AF488 and intrafoci distance histograms generated. In both cases, these histograms could be approximated with a Gaussian centered at an intrafoci distance of 135 nm with a width at half height of 70-80 nm. In cases where protein intrafoci distance histograms could similarly be fit with a Gaussian centered with the range 135 ± 37.5 nm, the relationship was considered intimate. In contrast, intrafoci distances which were approximated with a single Gaussian centered further than 180 nm described proteins occupying the same DSB but spatially separated.

Histograms not easily described by a single Gaussian were well approximated by fitting one Gaussian at 135 nm distance and 75 nm full width at half height with free fitting of the total area of the Gaussian, and then free fitting a second Gaussian to describe the remaining intrafoci distances. Fitting two Gaussians described arrangements where in some foci the proteins were closely associated while in others they were spatially separated. To further visualize the spread of intrafoci distances observed, the fitted Gaussians were extrapolated into 3D contoured heat maps by fitting a second Gaussian perpendicular to the 2D Gaussian distance/intensity coordinates with the area of the perpendicular Gaussian proportional to the intensity value of the fitted Gaussians. These contoured heatmaps, shown throughout in blue, describe the likelihood of particular intrafoci distances being observed for different protein combinations at naDNA.

All experiments were carried out in duplicate or more with repeats performed to generate approximately normal data distributions with N sizes not predetermined. Unequal variances, particularly across temporal series, are expected. Manual ROI selection of nuclei for all quantification, including data assessment for pairwise interdependency assessment and intrafoci association distribution was carried out blinded to the drug condition, protein species and time point being quantified. The representative images of cell nuclei and foci shown throughout were rendered at 10 nm, smoothed, dilated, and brightened to make them easily assessable after resizing for inclusion in figures. Importantly, all quantitative analysis used here was applied to images that had not been manually processed, removing any user bias. For examples of SR data processing and rendering for both analysis and user-friendly presentation within the paper see SI Figure S5.

### Comet Assays

A neutral comet assay was used to specifically assess the amount of DNA DSBs in cells treated with Veliparib, CPT, or both. In individual wells of a six-well plate, cells were incubated with 0.4 mL of Trypsin for 10 minutes. 1 mL of medium was added to the trypsin and the cells centrifuged at 1000 g for 5 minutes. The supernatant was removed, and the cell pellet was resuspended in 500 μL of PBS. The suspended cells were added to prewarmed low-melting agarose at 37°C (10 μL of cells to 90 μL of agarose). The mix was pipetted onto a CometSlide (4250, Trevigen) and spread equally across the slide before being allowed to set for 30 minutes at 4°C. Following this, the slides were submerged in cold Lysis Buffer (4250, Trevigen) for 30 minutes on a shaker, and then in cold Tris/Borate/EDTA buffer. Electrophoresis was run for 15 minutes at 21V in a Mini-Sub Cell GT electrophoresis tray (Bio-Rad). Subsequently, cells were fixed in 70% ethanol and allowed to dry overnight. DNA was stained with Cygreen (GEN-105, ENZO) for 30 minutes, and slides were imaged on an EVOS fluorescence microscope (AMG) with appropriate filters for GFP imaging. Images acquired were processed using Open Comet software and the olive moment for each group of cells was calculated [85].

### Statistics

Statistical analysis was carried out in OriginLab (8.5). All naDNA/protein overlap colocalization factors calculated for controls and CPT-treated cells were tested against control samples (Student’s two sample t-test). Combined Veliparib and CPT treatment was compared with controls, and with CPT-only samples, again using t-tests where appropriate.

### Code and data availability

The custom-written IDL mapping code, SR localization and rendering code, and the overlap analysis and Monte Carlo randomized simulation macros are available upon request. The data that support the findings of this study are available from the authors on reasonable request.

### Supplementary data statement

Supplementary data are available.

## Acknowledgements

We thank members of the Rothenberg laboratory for critically reading and commenting on the manuscript. Research in the Rothenberg laboratory is funded by grants from the National Institutes of Health (NIH: R01 GM108119), American Cancer Society (ACS: 130304-RSG-16-241-01-DMC), and the V Foundation for Cancer Research (D2018-020). D.R.W. is supported by a Bruce Stone Fellowship from the La Trobe Institute for Molecular Science and an Australian Research Council Discovery Early Career Research Award, and acknowledges funding from the Bendigo Tertiary Education Anniversary Foundation.

## Author Contributions

D.R.W and E.R conceived experiments, interpreted data, and wrote the manuscript with input from all authors. D.R.W, W.T.C.L, F.M and Y.T.K conducted the experiments. Y.Y provided analysis tools and expertise.

## References

1. Mehta A, Haber JE. Sources of DNA Double-Strand Breaks and Models of Recombinational DNA Repair. Cold Spring Harbor Perspectives in Biology. 2014;6(9). doi: 10.1101/cshperspect.a016428 PubMed PMID: WOS:000341576800008.

2. Ceccaldi R, Rondinelli B, D’Andrea AD. Repair Pathway Choices and Consequences at the Double-Strand Break. Trends in Cell Biology. 2016;26(1):52–64. doi: 10.1016/j.tcb.2015.07.009 PubMed PMID: WOS:000368206000007.

3. Neelsen KJ, Chaudhuri AR, Follonier C, Herrador R, Lopes M. Visualization and Interpretation of Eukaryotic DNA Replication Intermediates In Vivo by Electron Microscopy. Functional Analysis of DNA and Chromatin. 2014;1094:177–208. doi: 10.1007/978-1-62703-706-8_15. PubMed PMID: WOS:000332345200016.

4. Neelsen KJ, Lopes M. Replication fork reversal in eukaryotes: from dead end to dynamic response. Nature Reviews Molecular Cell Biology. 2015;16(4):207–20. doi: 10.1038/nrm3935 PubMed PMID: WOS:000351630500006.

5. Saleh-Gohari N, Bryant HE, Schultz N, Parker KA, Cassel TN, Helleday T. Spontaneous homologous recombination is induced by collapsed replication forks that are caused by endogenous DNA single-strand breaks. Molecular and Cellular Biology. 2005;25(16):7158–69. doi: 10.1128/mcb.25.16.7158-7169.2005 PubMed PMID: WOS:000231000800026.

6. Chaudhuri AR, Hashimoto Y, Herrador R, Neelsen KJ, Fachinetti D, Bermejo R, et al. Topoisomerase I poisoning results in PARP-mediated replication fork reversal. Nature Structural & Molecular Biology. 2012;19(4):417–23. doi: 10.1038/nsmb.2258 PubMed PMID: WOS:000302514400009.

7. Yeeles JTP, Poli J, Marians KJ, Pasero P. Rescuing stalled or damaged replication forks. Cold Spring Harbor perspectives in biology. 5(5):a012815-a. doi: 10.1101/cshperspect.a012815. PubMed PMID: 23637285.

8. Kolinjivadi AM, Sannino V, De Antoni A, Zadorozhny K, Kilkenny M, Techer H, et al. Smarcal1-Mediated Fork Reversal Triggers Mre11-Dependent Degradation of Nascent DNA in the Absence of Brca2 and Stable Rad51 Nucleofilaments. Molecular Cell. 2017;67(5):867-+. doi: 10.1016/j.molcel.2017.07.001. PubMed PMID: WOS:000411128900014.

9. Petermann E, Orta ML, Issaeva N, Schultz N, Helleday T. Hydroxyurea-Stalled Replication Forks Become Progressively Inactivated and Require Two Different RAD51-Mediated Pathways for Restart and Repair. Molecular Cell. 2010;37(4):492–502. doi: 10.1016/j.molcel.2010.01.021 PubMed PMID: PMC2958316.

10. Mazouzi A, Velimezi G, Loizou JI. DNA replication stress: Causes, resolution and disease. Exp Cell Res. 2014;329(1):85–93. doi: 10.1016/j.yexcr.2014.09.030 PubMed PMID: WOS:000345489700012.

11. Krejci L, Altmannova V, Spirek M, Zhao XL. Homologous recombination and its regulation. Nucleic Acids Res. 2012;40(13):5795–818. doi: 10.1093/nar/gks270 PubMed PMID: WOS:000306970700009.

12. Vilenchik MM, Knudson AG. Endogenous DNA double-strand breaks: Production, fidelity of repair, and induction of cancer. Proc Natl Acad Sci U S A. 2003;100(22):12871–6. doi: 10.1073/pnas.2135498100 PubMed PMID: WOS:000186301100066.

13. Heyer W-D, Ehmsen KT, Liu J. Regulation of Homologous Recombination in Eukaryotes. Annual Review of Genetics, Vol 44. 2010;44:113–39. doi: 10.1146/annurev-genet-051710-150955. PubMed PMID: WOS:000286042600006.

14. Willis NA, Chandramouly G, Huang B, Kwok A, Follonier C, Deng CX, et al. BRCA1 controls homologous recombination at Tus/Ter-stalled mammalian replication forks. Nature. 2014;510(7506):556-+. doi: 10.1038/nature13295. PubMed PMID: WOS:000337806300051.

15. Prakash R, Zhang Y, Feng WR, Jasin M. Homologous Recombination and Human Health: The Roles of BRCA1, BRCA2, and Associated Proteins. Cold Spring Harbor Perspectives in Biology. 2015;7(4). doi: 10.1101/cshperspect.a016600. PubMed PMID: WOS:000355194500006.

16. Liu T, Huang J. DNA End Resection: Facts and Mechanisms. Genomics Proteomics & Bioinformatics. 2016;14(3):126–30. doi: 10.1016/j.gpb.2016.05.002 PubMed PMID: WOS:000379516500003.

17. Daddacha W, Koyen AE, Bastien AJ, Head PE, Dhere VR, Nabeta GN, et al. SAMHD1 Promotes DNA End Resection to Facilitate DNA Repair by Homologous Recombination. Cell Reports. 2017;20(8):1921–35. doi: 10.1016/j.celrep.2017.08.008.

18. Whelan DR, Lee WTC, Yin Y, Ofri DM, Bermudez-Hernandez K, Keegan S, et al. Spatiotemporal dynamics of homologous recombination repair at single collapsed replication forks. Nature Communications. 2018;9(1):3882. doi: 10.1038/s41467-018-06435-3.

19. Schlacher K, Christ N, Siaud N, Egashira A, Wu H, Jasin M. Double-Strand Break Repair-Independent Role for BRCA2 in Blocking Stalled Replication Fork Degradation by MRE11. Cell. 2011;145(4):529–42. doi: 10.1016/j.cell.2011.03.041 PubMed PMID: WOS:000290560800007.

20. Ying S, Hamdy FC, Helleday T. Mre11-Dependent Degradation of Stalled DNA Replication Forks Is Prevented by BRCA2 and PARP1. Cancer Res. 2012;72(11):2814–21. doi: 10.1158/0008-5472.can-11-3417. PubMed PMID: WOS:000307348000014.

21. Chaudhuri AR, Callen E, Ding X, Gogola E, Duarte AA, Lee JE, et al. Replication fork stability confers chemoresistance in BRCA-deficient cells. Nature. 2016;535(7612):382-+. doi: 10.1038/nature18325. PubMed PMID: WOS:000380344200032.

22. Kolinjivadi AM, Sannino V, de Antoni A, Techer H, Baldi G, Costanzo V. Moonlighting at replication forks - a new life for homologous recombination proteins BRCA1, BRCA2 and RAD51. Febs Letters. 2017;591(8):1083-100. doi: 10.1002/1873-3468.12556. PubMed PMID: WOS:000400968800002.

23. Chapman JR, Taylor MRG, Boulton SJ. Playing the End Game: DNA Double-Strand Break Repair Pathway Choice. Molecular Cell. 2012;47(4):497–510. doi: 10.1016/j.molcel.2012.07.029 PubMed PMID: WOS:000308061100003.

24. Brandsma I, Gent DC. Pathway choice in DNA double strand break repair: observations of a balancing act. Genome Integrity. 2012;3:9-. doi: 10.1186/2041-9414-3-9. PubMed PMID: PMC3557175.

25. Arnoult N, Correia A, Ma J, Merlo A, Garcia-Gomez S, Maric M, et al. Regulation of DNA repair pathway choice in S and G2 phases by the NHEJ inhibitor CYREN. Nature. 2017;549(7673):548-+. doi: 10.1038/nature24023. PubMed PMID: WOS:000411930000056.

26. Hartlerode AJ, Morgan MJ, Wu Y, Buis J, Ferguson DO. Recruitment and activation of the ATM kinase in the absence of DNA-damage sensors. Nature Structural & Molecular Biology. 2015;22(9):736–U124. doi: 10.1038/nsmb.3072 PubMed PMID: WOS:000360933200017.

27. Lamarche BJ, Orazio NI, Weitzman MD. The MRN complex in Double-Strand Break Repair and Telomere Maintenance. FEBS letters. 2010;584(17):3682–95. doi: 10.1016/j.febslet.2010.07.029 PubMed PMID: PMC2946096.

28. Filippo JS, Sung P, Klein H. Mechanism of eukaryotic homologous recombination. Annual Review of Biochemistry. 2008;77:229–57. doi: 10.1146/annurev.biochern.77.061306.125255 PubMed PMID: WOS:000257596800011.

29. Sun J, Lee KJ, Davis AJ, Chen DJ. Human Ku70/80 Protein Blocks Exonuclease 1-mediated DNA Resection in the Presence of Human Mre11 or Mre11/Rad50 Protein Complex. J Biol Chem. 2012;287(7):4936–45. doi: 10.1074/jbc.M111.306167 PubMed PMID: WOS:000300608500054.

30. Lemacon D, Jackson J, Quinet A, Brickner JR, Li S, Yazinski S, et al. MRE11 and EXO1 nucleases degrade reversed forks and elicit MUS81-dependent fork rescue in BRCA2-deficient cells. Nature Communications. 2017;8. doi: 10.1038/s41467-017-01180-5 PubMed PMID: WOS:000412998100002.

31. Bryant HE, Petermann E, Schultz N, Jemth AS, Loseva O, Issaeva N, et al. PARP is activated at stalled forks to mediate Mre11-dependent replication restart and recombination. Embo Journal. 2009;28(17):2601–15. doi: 10.1038/emboj.2009.206 PubMed PMID: WOS:000269494200010.

32. Whelan DR, Bell TDM. Super-Resolution Single-Molecule Localization Microscopy: Tricks of the Trade. Journal of Physical Chemistry Letters. 2015;6(3):374–82. doi: 10.1021/jz5019702 PubMed PMID: WOS:000349137400012.

33. Raderschall E, Golub EI, Haaf T. Nuclear foci of mammalian recombination proteins are located at single-stranded DNA regions formed after DNA damage. Proc Natl Acad Sci U S A. 1999;96(5):1921–6. doi: 10.1073/pnas.96.5.1921 PubMed PMID: WOS:000078956600023.

34. Klein T, Proppert S, Sauer M. Eight years of single-molecule localization microscopy. Histochemistry and Cell Biology. 2014;141(6):561–75. doi: 10.1007/s00418-014-1184-3 PubMed PMID: WOS:000336388300002.

35. Whelan DR, Holm T, Sauer M, Bell TDM. Focus on Super-Resolution Imaging with Direct Stochastic Optical Reconstruction Microscopy (dSTORM). Australian Journal of Chemistry. 2014;67(2):179–83. doi: 10.1071/ch13499 PubMed PMID: WOS:000335560100001.

36. Heilemann M, van de Linde S, Schuttpelz M, Kasper R, Seefeldt B, Mukherjee A, et al. Subdiffraction-resolution fluorescence imaging with conventional fluorescent probes. Angewandte Chemie-International Edition. 2008;47(33):6172–6. doi: 10.1002/anie.200802376 PubMed PMID: WOS:000258355200005.

37. Reid DA, Keegan S, Leo-Macias A, Watanabe G, Strande NT, Chang HH, et al. Organization and dynamics of the nonhomologous end-joining machinery during DNA double-strand break repair. Proc Natl Acad Sci U S A. 2015;112(20):E2575–E84. doi: 10.1073/pnas.1420115112. PubMed PMID: WOS:000354729500007.

38. Zessin PJM, Finan K, Heilemann M. Super-resolution fluorescence imaging of chromosomal DNA. Journal of Structural Biology. 2012;177(2):344–8. doi: 10.1016/j.jsb.2011.12.015 PubMed PMID: WOS:000300755400019.

39. Raulf A, Spahn CK, Zessin PJM, Finan K, Bernhardt S, Heckel A, et al. Click chemistry facilitates direct labelling and super-resolution imaging of nucleic acids and proteins. Rsc Advances. 2014;4(57):30462–6. doi: 10.1039/c4ra01027b PubMed PMID: WOS:000340497600069.

40. Chagin VO, Casas-Delucchi CS, Reinhart M, Schermelleh L, Markaki Y, Maiser A, et al. 4D Visualization of replication foci in mammalian cells corresponding to individual replicons. Nature Communications. 2016;7. doi: 10.1038/ncomms11231 PubMed PMID: WOS:000373827000001.

41. Triemer T, Messikommer A, Glasauer SMK, Alzeer J, Paulisch MH, Luedtke NW. Superresolution imaging of individual replication forks reveals unexpected prodrug resistance mechanism. Proceedings of the National Academy of Sciences. 2018;115(7):E1366. doi: 10.1073/pnas.1714790115.

42. Xiang W, Roberti MJ, Hériché J-K, Huet S, Alexander S, Ellenberg J. Correlative live and super-resolution imaging reveals the dynamic structure of replication domains. The Journal of Cell Biology. 2018;217(6):1973. doi: 10.1083/jcb.201709074.

43. Gorczyca W, Gong JP, Darzynkiewicz Z. DETECTION OF DNA STRAND BREAKS IN INDIVIDUAL APOPTOTIC CELLS BY THE INSITU TERMINAL DEOXYNUCLEOTIDYL TRANSFERASE AND NICK TRANSLATION ASSAYS. Cancer Res. 1993;53(8):1945–51. PubMed PMID: WOS:A1993KY09600040.

44. Mijic S, Zellweger R, Chappidi N, Berti M, Jacobs K, Mutreja K, et al. Replication fork reversal triggers fork degradation in BRCA2-defective cells. Nature Communications. 2017;8(1):859. doi: 10.1038/s41467-017-01164-5.

45. Zellweger R, Dalcher D, Mutreja K, Berti M, Schmid JA, Herrador R, et al. Rad51-mediated replication fork reversal is a global response to genotoxic treatments in human cells. Journal of Cell Biology. 2015;208(5):563–79. doi: 10.1083/jcb.201406099 PubMed PMID: WOS:000350530600009.

46. Berti M, Ray Chaudhuri A, Thangavel S, Gomathinayagam S, Kenig S, Vujanovic M, et al. Human RECQ1 promotes restart of replication forks reversed by DNA topoisomerase I inhibition. Nature Structural & Molecular Biology. 2013;20:347. doi: 10.1038/nsmb.2501 https://www.nature.com/articles/nsmb.2501#supplementary-information.

47. Chanut P, Britton S, Coates J, Jackson SP, Calsou P. Coordinated nuclease activities counteract Ku at single-ended DNA double-strand breaks. Nature Communications. 2016;7:12889. doi: 10.1038/ncomms12889 https://www.nature.com/articles/ncomms12889#supplementary-information.

48. Britton S, Coates J, Jackson SP. A new method for high-resolution imaging of Ku foci to decipher mechanisms of DNA double-strand break repair. Journal of Cell Biology. 2013;202(3):579–95. doi: 10.1083/jcb.201303073 PubMed PMID: WOS:000322769400016.

49. Garcia V, Phelps SEL, Gray S, Neale MJ. Bidirectional resection of DNA double-strand breaks by Mre11 and Exo1. Nature. 2011;479(7372):241–U123. doi: 10.1038/nature10515. PubMed PMID: WOS:000298030800045.

50. Bermudez-Hernandez K, Keegan S, Whelan DR, Reid DA, Zagelbaum J, Yin Y, et al. A Method for Quantifying Molecular Interactions Using Stochastic Modelling and Super-Resolution Microscopy. Scientific Reports. 2017;7(1):14882. doi: 10.1038/s41598-017-14922-8.

51. MacPhail SH, Banath JP, Yu Y, Chu E, Olive PL. Cell cycle-dependent expression of phosphorylated histone H2AX: Reduced expression in unirradiated but not X-irradiated G(1)-phase cells. Radiat Res. 2003;159(6):759–67. doi: 10.1667/rr3003 PubMed PMID: WOS:000183291800008.

52. Bianchi A, de Lange T. Ku binds telomeric DNA in vitro. J Biol Chem. 1999;274(30):21223–7. doi: 10.1074/jbc.274.30.21223 PubMed PMID: WOS:000081613100069.

53. de Jager M, Dronkert MLG, Modesti M, Beerens C, Kanaar R, van Gent DC. DNA-binding and strand-annealing activities of human Mre11: implications for its roles in DNA double-strand break repair pathways. Nucleic Acids Res. 2001;29(6):1317–25. doi: 10.1093/nar/29.6.1317 PubMed PMID: WOS:000167529200008.

54. Davis AJ, Chen DJ. DNA double strand break repair via non-homologous end-joining. Translational cancer research. 2013;2(3):130–43. doi: 10.3978/j.issn.2218-676X.2013.04.02 PubMed PMID: PMC3758668.

55. Daley JM, Sung P. 53BP1, BRCA1, and the Choice between Recombination and End Joining at DNA Double-Strand Breaks. Molecular and Cellular Biology. 2014;34(8):1380–8. doi: 10.1128/MCB.01639-13. PubMed PMID: PMC3993578.

56. Chung W-H, Zhu Z, Papusha A, Malkova A, Ira G. Defective Resection at DNA Double-Strand Breaks Leads to De Novo Telomere Formation and Enhances Gene Targeting. PLoS Genetics. 2010;6(5):e1000948. doi: 10.1371/journal.pgen.1000948 PubMed PMID: PMC2869328.

57. Shiloh Y, Schans GPVD, Lohman PHML, Becker Y. Induction and repair of DNA damage in normal and ataxiatelangiectasia skin fibroblasts treated with neocarzinostatin. Carcinogenesis. 1983;4(7):917–21. doi: 10.1093/carcin/4.7.917.

58. Lee C-S, Lee K, Legube G, Haber JE. Dynamics of yeast histone H2A and H2B phosphorylation in response to a double-strand break. Nature Structural & Molecular Biology. 2013;21:103. doi: 10.1038/nsmb.2737 https://www.nature.com/articles/nsmb.2737#supplementary-information.

59. Mah L-J, El-Osta A, Karagiannis TC. γH2AX as a molecular marker of aging and disease. Epigenetics. 2010;5(2):129–36. doi: 10.4161/epi.5.2.11080.

60. Shao Z, Davis AJ, Fattah KR, So S, Sun J, Lee K-J, et al. Persistently bound Ku at DNA ends attenuates DNA end resection and homologous recombination. DNA Repair. 2012;11(3):310–6. doi: 10.1016/j.dnarep.2011.12.007 PubMed PMID: PMC3297478.

61. Lee K-J, Saha J, Sun J, Fattah KR, Wang S-C, Jakob B, et al. Phosphorylation of Ku dictates DNA double-strand break (DSB) repair pathway choice in S phase. Nucleic Acids Res. 2016;44(4):1732–45. doi: 10.1093/nar/gkv1499.

62. Andrews BJ, Lehman JA, Turchi JJ. Kinetic analysis of the Ku-DNA binding activity reveals a redox-dependent alteration in protein structure that stimulates dissociation of the Ku-DNA complex. J Biol Chem. 2006;281(19):13596–603. doi: 10.1074/jbc.M512787200 PubMed PMID: WOS:000237336600069.

63. Postow L, Ghenoiu C, Woo EM, Krutchinsky AN, Chait BT, Funabiki H. Ku80 removal from DNA through double strand break-induced ubiquitylation. Journal of Cell Biology. 2008;182(3):467–79. doi: 10.1083/jcb.200802146 PubMed PMID: WOS:000258529100008.

64. Krasner DS, Daley JM, Sung P, Niu HY. Interplay between Ku and Replication Protein A in the Restriction of Exo1-mediated DNA Break End Resection. J Biol Chem. 2015;290(30):18806–16. doi: 10.1074/jbc.M115.660191 PubMed PMID: WOS:000358512100047.

65. Bakr A, Oing C, Kocher S, Borgmann K, Dornreiter I, Petersen C, et al. Involvement of ATM in homologous recombination after end resection and RAD51 nucleofilament formation. Nucleic Acids Res. 2015;43(6):3154–66. doi: 10.1093/nar/gkv160 PubMed PMID: WOS:000354719300022.

66. Godthelp BC, Artwert F, Joenje H, Zdzienicka MZ. Impaired DNA damage-induced nuclear Rad51 foci formation uniquely characterizes Fanconi anemia group D1. Oncogene. 2002;21(32):5002–5. doi: 10.1038/sj.onc.1205656 PubMed PMID: WOS:000176874800016.

67. Paull TT, Rogakou EP, Yamazaki V, Kirchgessner CU, Gellert M, Bonner WM. A critical role for histone H2AX in recruitment of repair factors to nuclear foci after DNA damage. Current Biology. 2000;10(15):886–95. doi: 10.1016/s0960-9822(00)00610-2 PubMed PMID: WOS:000088979300016.

68. Wray J, Liu JM, Nickoloff JA, Shen ZY. Distinct RAD51 associations with RAD52 and BCCIP in response to DNA damage and replication stress. Cancer Res. 2008;68(8):2699–707. doi: 10.1158/0008-5472.can-07-6505 PubMed PMID: WOS:000255100500020.

69. Liu YL, Maizels N. Coordinated response of mammalian Rad51 and Rad52 to DNA damage. Embo Reports. 2000;1(1):85–90. doi: 10.1093/emborep.kvd002 PubMed PMID: WOS:000165765800020.

70. Li L. BRCA1 Forks Over New Roles in DNA-Damage Response– Before and Beyond the Breaks. Molecular Cell. 2011;44(2):174–6. doi: https://doi.org/10.1016/j.molcel.2011.10.003.

71. Gonzalez-Prieto R, Munoz-Cabello AM, Cabello-Lobato MJ, Prado F. Rad51 replication fork recruitment is required for DNA damage tolerance. Embo Journal. 2013;32(9):1307–21. doi: 10.1038/emboj.2013.73 PubMed PMID: WOS:000319122600011.

72. Irmisch A, Ampatzidou E, Mizuno K, O’Connell MJ, Murray JM. Smc5/6 maintains stalled replication forks in a recombination-competent conformation. Embo Journal. 2009;28(2):144–55. doi: 10.1038/emboj.2008.273 PubMed PMID: WOS:000262580800009.

73. Hanamshet K, Mazina OM, Mazin AV. Reappearance from Obscurity: Mammalian Rad52 in Homologous Recombination. Genes. 2016;7(9). doi: 10.3390/genes7090063 PubMed PMID: WOS:000385535300009.

74. Lok BH, Carley AC, Tchang B, Powell SN. RAD52 inactivation is synthetically lethal with deficiencies in BRCA1 and PALB2 in addition to BRCA2 through RAD51-mediated homologous recombination. Oncogene. 2013;32(30):3552–8. doi: 10.1038/onc.2012.391 PubMed PMID: WOS:000322220800008.

75. Maya-Mendoza A, Moudry P, Merchut-Maya JM, Lee M, Strauss R, Bartek J. High speed of fork progression induces DNA replication stress and genomic instability. Nature. 2018;559(7713):279–84. doi: 10.1038/s41586-018-0261-5.

76. Karanam K, Kafri R, Loewer A, Lahav G. Quantitative Live Cell Imaging Reveals a Gradual Shift between DNA Repair Mechanisms and a Maximal Use of HR in Mid S Phase. Molecular Cell. 2012;47(2):320–9. doi: 10.1016/j.molcel.2012.05.052 PubMed PMID: WOS:000307084000017.

77. Myler LR, Gallardo IF, Soniat MM, Deshpande RA, Gonzalez XB, Kim Y, et al. Single-Molecule Imaging Reveals How Mre11-Rad50-Nbs1 Initiates DNA Break Repair. Molecular Cell. 2017;67(5):891–8.e4. doi: 10.1016/j.molcel.2017.08.002.

78. Chen Y-H, Jones MJK, Yin Y, Crist SB, Colnaghi L, Sims RJ, III, et al. ATR-Mediated Phosphorylation of FANCI Regulates Dormant Origin Firing in Response to Replication Stress. Molecular Cell. 2015;58(2):323–38. doi: 10.1016/j.molcel.2015.02.031 PubMed PMID: WOS:000353222900013.

79. Yin YD, Rothenberg E. Probing the Spatial Organization of Molecular Complexes Using Triple-Pair-Correlation. Scientific Reports. 2016;6. doi: 10.1038/srep30819 PubMed PMID: WOS:000381698000001.

80. Cooper S. Reappraisal of serum starvation, the restriction point, G0, and G1 phase arrest points. Faseb Journal. 2003;17(3):333–40. doi: 10.1096/fj.02-0352rev. PubMed PMID: WOS:000181892600032.

81. Wagner LM. Profile of veliparib and its potential in the treatment of solid tumors. Oncotargets and Therapy. 2015;8:1931–9. doi: 10.2147/ott.s69935 PubMed PMID: WOS:000358650600001.

82. Whelan DR, Bell TDM. Image artifacts in Single Molecule Localization Microscopy: why optimization of sample preparation protocols matters. Scientific Reports. 2015;5. doi: 10.1038/srep07924 PubMed PMID: WOS:000348104600002.

83. Schneider CA, Rasband WS, Eliceiri KW. NIH Image to ImageJ: 25 years of image analysis. Nature Methods. 2012;9(7):671–5. doi: 10.1038/nmeth.2089 PubMed PMID: WOS:000305942200020.

84. Henriques R, Lelek M, Fornasiero EF, Valtorta F, Zimmer C, Mhlanga MM. QuickPALM: 3D real-time photoactivation nanoscopy image processing in ImageJ. Nature Methods. 2010;7(5):339–40. doi: 10.1038/nmeth0510-339 PubMed PMID: WOS:000277175100006.

85. Gyori BM, Venkatachalam G, Thiagarajan PS, Hsu D, Clement MV. Open Comet: An automated tool for comet assay image analysis. Redox Biology. 2014;2:457–65. doi: 10.1016/j.rcdox.2013.12.020 PubMed PMID: WOS:000350769600055.

