## Supplemental Data for "Super-resolution visualization of distinct stalled and broken replication fork structures"

**Contents.**

**Supplementary Figures 1-5**
**Supplementary Tables 1-3**

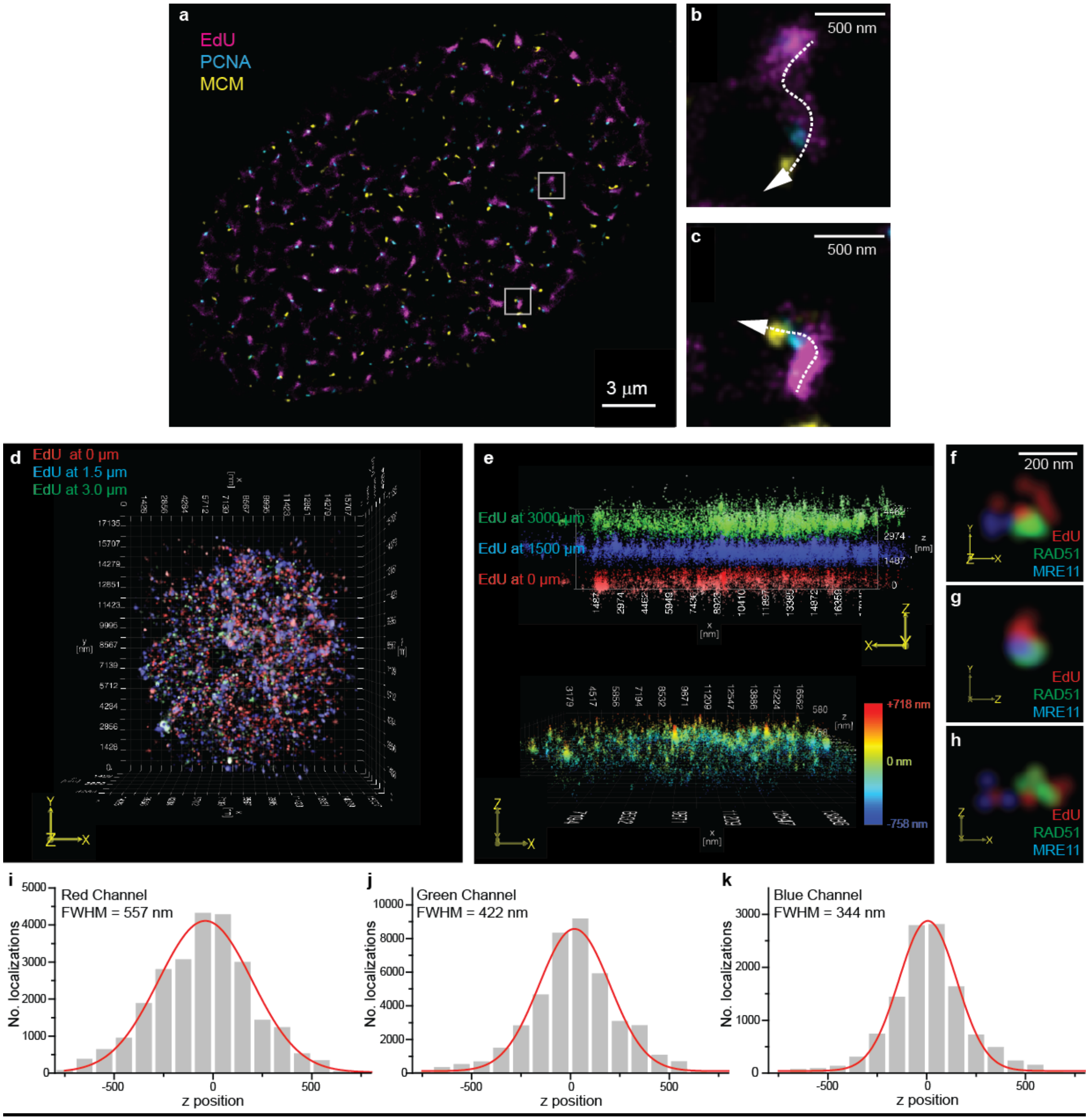

Figure S1

Individual repair foci comprise an individual seDSB and the 3D organization can be interpreted using 2D multi-color super resolution images.

(A) To confirm the individual nature of single RFs within single foci we prepared cells in mid-S phase pulse labeled with EdU and immunlabeled for PCNA (PC10, Abcam) and MCM (EP2863Y, Abcam).

(B-C) Elongated replicons were the prevalent species throughout the cell and showed the expected sequential nature of MCM-PCNA-naDNA confirming the RF was progressed as a single entity and not in a 'factory'. In conjunction with our regular detection of only single protein foci for any one naDNA we therefore concluded that we could assess naDNA foci as typically containing no, or one, damaged RF.

(D-E) Cells pulse labeled with EdU were imaged in 3D to assess the typical axial depth of images contained using our highly inclined laminated optical sheet 2D acquisition setup. By moving the focus axially from a low focal plane (starting at 0 µm which we judged to be near the bottom of the nucleus) to two higher planes (+1.5 and +3.0 µm) and then resolving the EdU distribution in 3D we demonstrate that with HiLo illumination was are only sampling a ~1 µm thick slice of the cell, excluding interference from out of plane EdU. A representative cell is shown in 3D in the xy plane (D) and the xz plane (E, upper) with each of the three measurement planes depicted in a different color. E, lower, shows a single 3D projection in xz space demonstrating that the majority of detected molecules exist ±500 nm from the imaging plane.

(F-H) Costaining of EdU, BRCA2, and RAD51 8 hours after CPT damage resulted in the characteristic 3D images of small repair foci showing all three colors. While different perspectives offered interesting insights into the intrafoci arrangement, colocalization was visible from all angles.

(I-K) Quantification of the axial position of detected localizations in z-space for red, green, and blue labeled samples show axial sampling depths of 344 nm in blue, 422 nm in green, and 557 nm in red due to the chromatic differences in the emissions.

**Analysis Description** | **Example** | **Characteristics**

**A** **(i)** Two-color analysis of the percentage of naDNA foci overlapped with a specific protein as compared with control and random levels of overlap. **(ii)** The generated heatmap normalized to control levels and depicting significance.

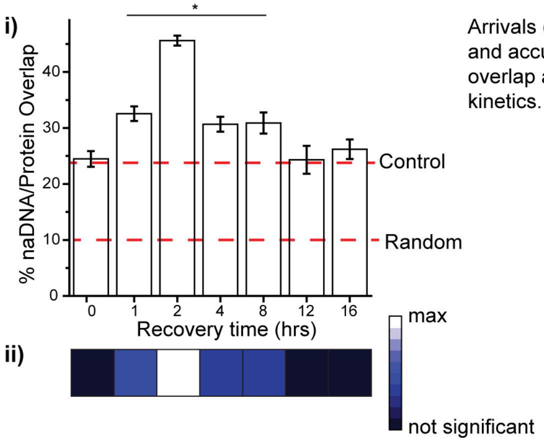

Arrivals (percentage overlap count) and accumulations (percentage overlap area) providing overall kinetics.

**B** Pairwise quantification of the proportion of naDNA foci positive for one or both proteins stained.

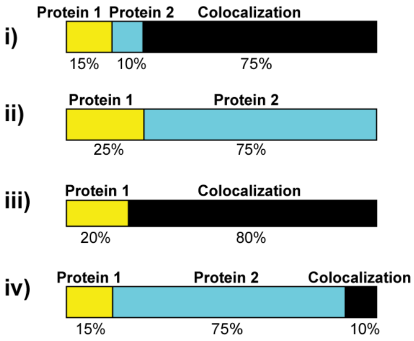

Predominance of **(i)** colocalization, **(ii)** exclusivity, **(iii)** dependence, or **(iv)** the prevalence of one protein over another.

**C** Analysis of three color foci (two species colocalized with naDNA) to determine the rendered SR distance between the two proteins at the single DSB site. Conversion of the histogrammed frequency of inter-species distances (grey) allows generation of a 2D distribution map (blue).

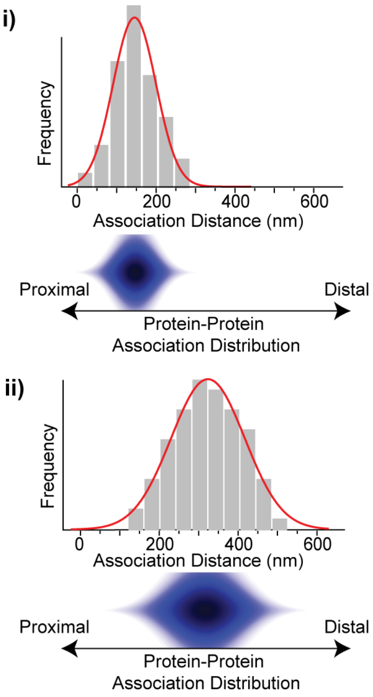

Description of protein-protein association as either **(i)** proximal or **(ii)** distal which allows identification of DSB-associated proteins as either complexed/interacting or spatially separated/independent, respectively.

Figure S2

Three-tiered analytical approach for determining the overall kinetics of HR, the colocalization of proteins at single foci, and the internal organization of these proteins. Related to Figures 1-4.

(A) A description, example, and principal characteristics elucidated by quantification of protein overlap with naDNA. Automated thresholding of SR images enabled pulse-labeled single RF naDNA foci to be defined and examined for overlap with ssDNA, TUNEL signal (DSBs), and various protein localizations. The total number of overlaps per cell was found to sensitively describe the kinetics of proteins expected to interact with stressed RFs in low numbers, whereas the total area of overlaps better described proteins expected to accumulate, such as RAD51, BRCA2, and RPA. (i) The number or area of naDNA/protein overlap was normalized to the level of overlaps predicted using randomized Monte Carlo simulations to account for the dense nuclear environment, and to the level of overlap detected in undamaged cells.

(B) If proteins were found to significantly colocalize with naDNA after damage, they were further assessed in a pairwise fashion to quantify of the extent of co-occupancy at naDNA foci versus foci positive for only one or the other protein. In this way, we could determine the predominance of (i) colocalization, (ii) exclusivity, whereby the presence of one protein excluded the second protein from associating, (iii) dependence, whereby the presence of one protein was dependent on the presence of the other, or (iv) the prevalence of one protein over another.

(C) Finally, pairwise labeling of proteins that yielded high incidences of protein-protein colocalization with naDNA were examined to determine the internal organization of proteins within these single foci. To do this, the distance between the centers of mass of the fluorophore localization clusters attributed to each protein was measured. A histogram of the distances detected could then be transformed into a 3D protein-protein distribution map which depicted either a (i) proximal or (ii) distal relationship between the two proteins under examination. This allowed differentiation between proteins that are spatially close to each other and potentially interacting from those that are at the same repair foci but are spatially separated and not interacting.

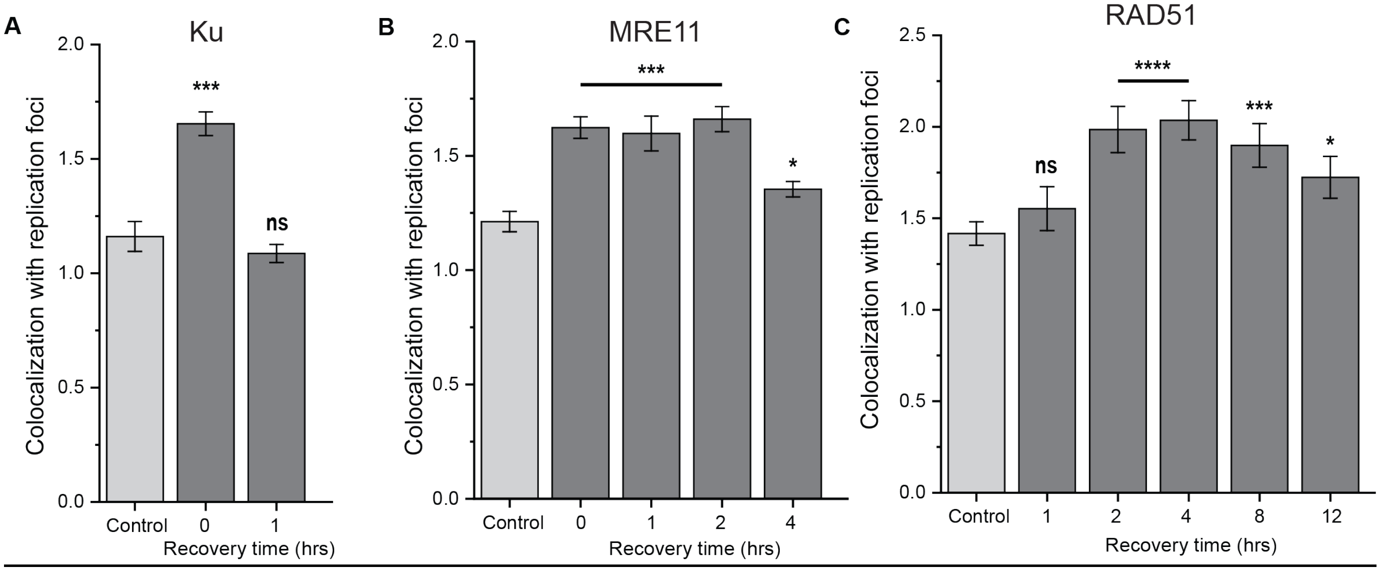

Figure S3

Temporal spatiokinetics of Ku, MRE11, and RAD51 at seDSBs.

(A) Quantification of colocalization of Ku with naDNA foci in control cells and 0 and 1 hour after CPT-treatment showing Ku association only at 0 hours.

(B) Quantification of colocalization of MRE11 with naDNA foci in control cells and 0, 1, 2 and 4 hours after CPT-treatment showing MRE11 association 0-2 hours indicating ongoing resection.

(C) Quantification of colocalization of RAD51 with naDNA foci in control cells and 1, 2, 4, 8, and 12 hours after CPT-treatment showing no RAD51 association at 1 hour, peak RAD51 association at 2-4 hours before dissipation. This indicates the kinetics of RAD51/ssDNA nucleoprotein filament formation and homology search.

Complete N values available in Tables S1-2. All graphs show mean ± s.e.m. Student's t-test results shown for comparison with control levels: ns depicts p>0.05, * depicts p<0.05, ** depicts p<0.01, *** depicts p<0.001, **** depicts p<0.0001.

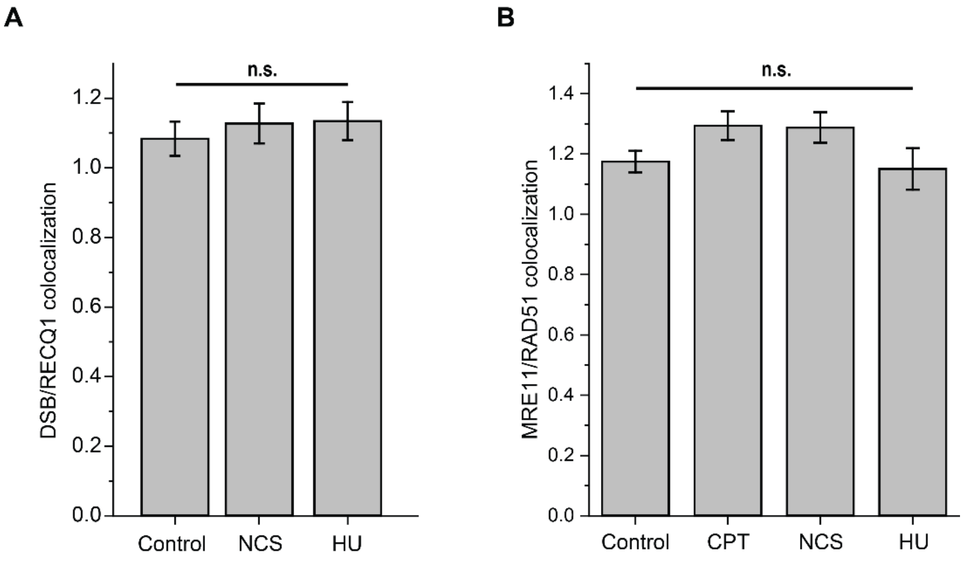

Figure S4

Colocalization of DSBs and RECQ1, and MRE11 and RAD51, in control cells.

(A) Quantification of colocalization of DSBs (TUNEL) with RECQ1 in control cells and cells damaged using NCS or HU demonstrating no significant association.

(B) Quantification of colocalization of MRE11 with RAD51 foci in control cells and in cells immediately following damage using CPT, NCS and HU.

Complete N values available in Tables S1-2**.** All graphs show mean ± s.e.m. Student's t-test results shown for comparison with control levels: ns depicts $p>0.05$

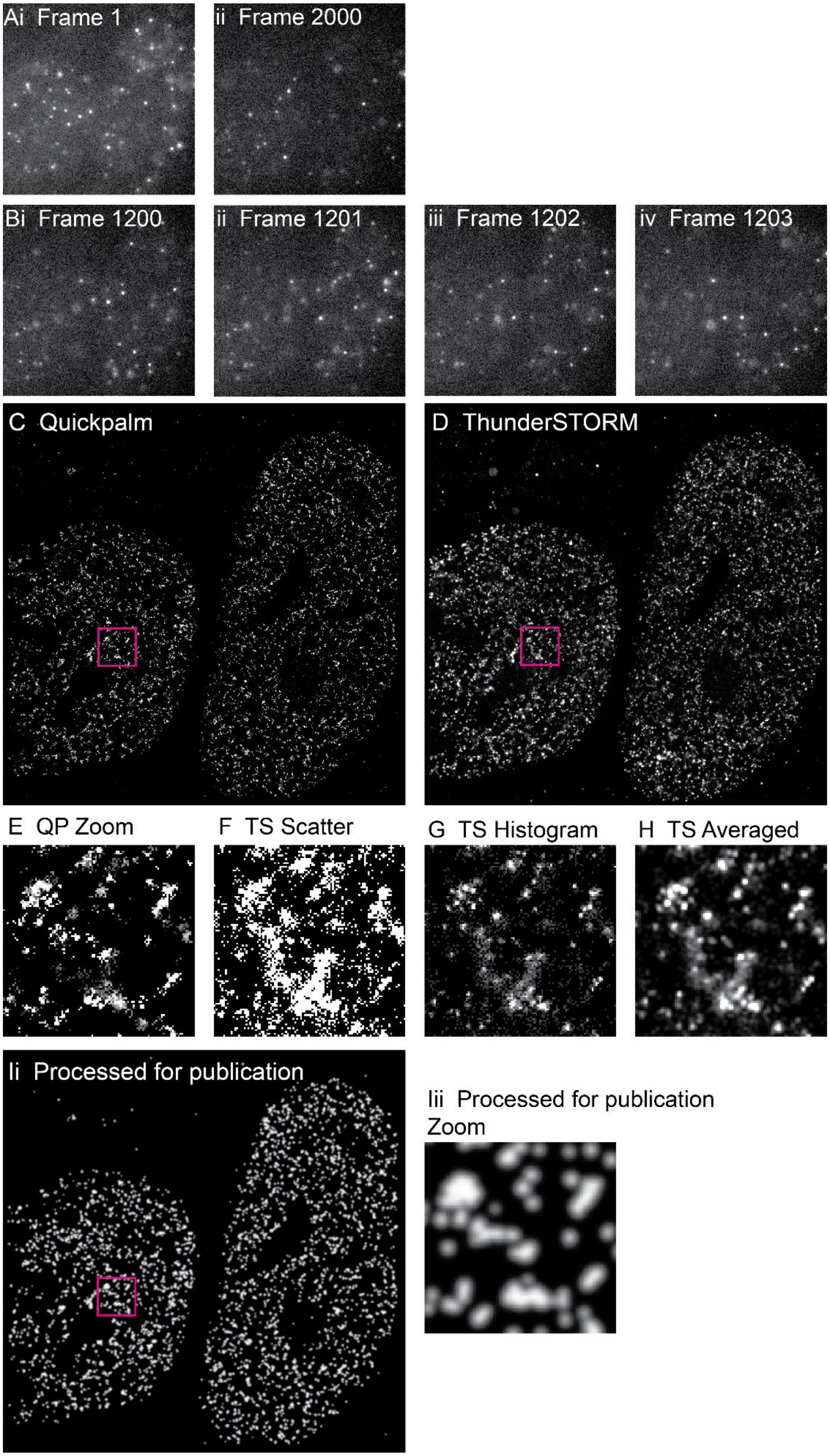

Ai Frame 1
ii Frame 2000
Bi Frame 1200
ii Frame 1201
iii Frame 1202
iv Frame 1203
C Quickpalm
D ThunderSTORM
E QP Zoom
F TS Scatter
G TS Histogram
H TS Averaged
Ii Processed for publication
Iii Processed for publication Zoom

Figure S5

Raw and processed SR data showing blinking and rendering quality.

(Ai) First and (ii) last frames in a representative raw image stack taken using red excitation/channel settings.

(Bi-iv) Four sequential frames taken from the raw image stack as in (A), representative of typical mid-movie blinking.

(C) The raw QuickPALM output image, brightened as a 32 bit image.

(D) The Averaged histograms ThunderSTORM output image.

(E) A zoom section from (C).

(F) A zoom section as in (C) showing the ThunderSTORM scatter plot output.

(G) A zoom section as in (C) showing the ThunderSTORM histogram plot output.

(H) A zoom section as in (C) from (D) showing the ThunderSTORM comparison.

(Ii) The QuickPALM image processed using smoothing and binarizing algorithms for display purposes within the manuscript. Analysis was performed on raw outputs as shown in (D). (ii) The coinciding zoom area.

All zoomed out images including (A-B) are of ~25 μm across fields of view. All zoomed in frames are 2 μm across.

### Table S1: N values for overlap analyses.

| Drug Condition | Time | Species 1 | Species 2 | N |
|---|---|---|---|---|
| Control | 0 | naDNA | yH2A.X | 30 |
| CPT Only | 0 | naDNA | yH2A.X | 40 |
| Control | 0 | naDNA | TUNEL | 31 |
| CPT Only (Fig 1C) | 0 | naDNA | TUNEL | 82 |
| CPT Only | 0 | naDNA | TUNEL | 42 |
| CPT +Veliparib | 0 | naDNA | TUNEL | 62 |
| Control | 0 | naDNA | Ku | 19 |
| CPT Only (Fig 1C) | 0 | naDNA | Ku | 121 |
| CPT Only | 0 | naDNA | Ku | 103 |
| CPT Only | 1 | naDNA | Ku | 51 |
| CPT+Veliparib | 0 | naDNA | Ku | 42 |
| Control | 0 | naDNA | MRE11 | 37 |
| CPT Only (Fig 1C) | 0 | naDNA | MRE11 | 99 |
| CPT Only | 0 | naDNA | MRE11 | 96 |
| CPT Only | 1 | naDNA | MRE11 | 48 |
| CPT Only | 2 | naDNA | MRE11 | 104 |
| CPT Only | 4 | naDNA | MRE11 | 76 |
| CPT+Veliparib | 0 | naDNA | MRE11 | 46 |
| Control | 0 | naDNA | RAD52 | 18 |
| CPT Only | 0 | naDNA | RAD52 | 24 |
| CPT Only | 1 | naDNA | RAD52 | 13 |
| HU | 0 | naDNA | RAD52 | 25 |
| HU | 1.5 | naDNA | RAD52 | 31 |
| Control | 0 | RAD51 | RAD52 | 39 |
| CPT Only | 0 | RAD51 | RAD52 | 27 |
| CPT Only | 1 | RAD51 | RAD52 | 19 |

| Drug Condition | Time | Species 1 | Species 2 | N |
|---|---|---|---|---|
| Control | 0 | naDNA | BRCA1 | 46 |
| HU | 0 | naDNA | BRCA1 | 30 |
| HU | 1.5 | naDNA | BRCA1 | 30 |
| Control | 0 | naDNA | RAD51 | 67 |
| CPT Only | 0 | naDNA | RAD51 | 57 |
| CPT Only | 1 | naDNA | RAD51 | 49 |
| CPT Only | 2 | naDNA | RAD51 | 50 |
| CPT Only | 4 | naDNA | RAD51 | 47 |
| CPT Only | 8 | naDNA | RAD51 | 53 |
| CPT Only | 12 | naDNA | RAD51 | 42 |
| HU | 0 | naDNA | RAD51 | 34 |
| HU | 1.5 | naDNA | RAD51 | 39 |
| Control | 0 | naDNA | RECQ1 | 56 |
| CPT Only (Fig 1C) | 0 | naDNA | RECQ1 | 78 |
| CPT Only | 0 | naDNA | RECQ1 | 49 |
| Control | 0 | naDNA | RPA | 63 |
| HU | 0 | naDNA | RPA | 34 |
| HU | 1.5 | naDNA | RPA | 39 |
| Control | 0 | naDNA | TopI | 24 |
| CPT Only | 0 | naDNA | TopI | 35 |
| Control | 0 | DSB | RAD51 | 71 |
| NCS | 0 | DSB | RAD51 | 64 |
| Control | 0 | DSB | RAD52 | 72 |
| NCS | 0 | DSB | RAD52 | 45 |

| Drug Condition | Time | Species 1 | Species 2 | N |
|---|---|---|---|---|
| Control | 0 | DSB | Ku | 61 |
| NCS | 0 | DSB | Ku | 72 |
| HU | 0 | DSB | Ku | 51 |
| Control | 0 | DSB | MRE11 | 56 |
| NCS | 0 | DSB | MRE11 | 36 |
| HU | 0 | DSB | MRE11 | 70 |

**Table S2: N values for intrafoci analyses of WT+CPT damaged cells.**

| Time | Species 1 | Species 2 | N |
|---|---|---|---|
| 0 | MRE11 | Ku | 107 |
| 0 | TopI | DSB | 65 |
| 0 | MRE11 | DSB | 51 |
| 0 | Ku | DSB | 53 |
| 0 | RAD51 | DSB | 94 |
| 0 | Ku | RAD51 | 101 |
| 0 | MRE11 | RAD51 | 76 |
| 0 | RAD52 | RAD51 | 58 |
| 0 | Ku | RECQ1 | 145 |
| 0 | RAD51 | RECQ1 | 101 |

**Table S3: Antibody List.** Related to Figures 1-4.

| Target | Species/Conjugate | Product Code | Manufacturer | Dilutions | Refs |
|---|---|---|---|---|---|
| BRCA1 | mouse monoclonal AF488 conjugated | NB100-598AF488 | Novus | 1:250 | * |
| Ku | mouse monoclonal | ms-286 | ThermoFisher | 1:1000/1:5000 | 1 |
| MRE11 | rabbit polyclonal | NB100-142 | Novus | 1:500/1:2000 | 2, 3 |
| RAD51 | rabbit monoclonal AF488 conjugated | AB196449 | Abcam | 1:250 | * |
| RAD51 | rabbit polyclonal | 39194 | Active Motif | 1:500/1:1000 | 4 |
| RAD51 | mouse monoclonal | GTX70230 | Genetex | 1:500/1:2000 | 5 |
| RAD52 | rabbit polyclonal | SC8350 | Santa Cruz | 1:400/1:2000 | 6 |
| RECQ1 | rabbit polyclonal | Ab151501 | Abcam | 1:500/1:2000 | * |
| RECQ1 | mouse polyclonal | Ab89817 | Abcam | 1:200/1:1000 | * |
| RPA | mouse monoclonal | AB2175 | Abcam | 1:500/1:2000 | 7, 8 |
| RPA | rabbit monoclonal AF488 conjugated | AB199097 | Abcam | 1:300 | 9 |
| TopI | rabbit polyclonal | AB3825 | Abcam | 1:500/1:2000 | 10 |
| yH2A.X | rabbit polyclonal | NB100-384 | Novus | 1:2000/1:10000 | 11,12 |
| yH2A.X | mouse monoclonal | 05-636 | EMD Millipore | 1:2000/1:10000 | 13 |
| | goat-anti-rabbit AF568 | A11036 | Invitrogen | | |
| | goat-anti-rabbit AF488 | A11034 | Invitrogen | | |
| | goat-anti-mouse AF568 | A11031 | Invitrogen | | |
| | goat anti-mouse AF488 | A11029 | Invitrogen | | |

* denotes antibodies used which had not been used for IF applications in publications previously. To validate these antibodies they were double-stained alongside antibodies for the same target and found to have good colocalization.

1 Reid, D. A. *et al.* Organization and dynamics of the nonhomologous end-joining machinery during DNA double-strand break repair. *Proc. Natl. Acad. Sci. U. S. A.* **112**, E2575-E2584, doi:10.1073/pnas.1420115112 (2015).

2 Lee, K. Y. *et al.* MCM8-9 complex promotes resection of double-strand break ends by MRE11-RAD50-NBS1 complex. *Nature Communications* **6**, doi:10.1038/ncomms8744 (2015).

3 Gao, M. *et al.* Ago2 facilitates Rad51 recruitment and DNA double-strand break repair by homologous recombination. *Cell Research* **24**, 532-541, doi:10.1038/cr.2014.36 (2014).

4 Bennett, B. T. & Knight, K. L. Cellular localization of human Rad51C and regulation of ubiquitin-mediated proteolysis of Rad51. *J. Cell. Biochem.* **96**, 1095-1109, doi:10.1002/jcb.20640 (2005).

5 Pfaffle, H. N. *et al.* EGFR-Activating Mutations Correlate with a Fanconi Anemia-like Cellular Phenotype That Includes PARP Inhibitor Sensitivity. *Cancer Res.* **73**, 6254-6263, doi:10.1158/0008-5472.can-13-0044 (2013).

6 Wray, J., Liu, J. M., Nickoloff, J. A. & Shen, Z. Y. Distinct RAD51 associations with RAD52 and BCCIP in response to DNA damage and replication stress. *Cancer Res.* **68**, 2699-2707, doi:10.1158/0008-5472.can-07-6505 (2008).

7 Leung, J. W. *et al.* Nucleosome Acidic Patch Promotes RNF168-and RING1B/BMI1-Dependent H2AX and H2A Ubiquitination and DNA Damage Signaling. *Plos Genetics* **10**, doi:10.1371/journal.pgen.1004178 (2014).

8 Carvalho, S. *et al.* SETD2 is required for DNA double-strand break repair and activation of the p53-mediated checkpoint. *Elife* **3**, doi:10.7554/eLife.02482 (2014).

9 Toledo, L. I. *et al.* ATR Prohibits Replication Catastrophe by Preventing Global Exhaustion of RPA. *Cell* **155**, 1088-1103, doi:10.1016/j.cell.2013.10.043 (2013).

10 Ye, J. *et al.* TRF2 and Apollo Cooperate with Topoisomerase 2 alpha to Protect Human Telomeres from Replicative Damage. *Cell* **142**, 230-242, doi:10.1016/j.cell.2010.05.032 (2010).

11 Markova, E. *et al.* DNA repair foci and late apoptosis/necrosis in peripheral blood lymphocytes of breast cancer patients undergoing radiotherapy. *International Journal of Radiation Biology* **91**, 934-945, doi:10.3109/09553002.2015.1101498 (2015).

12 Francia, S., Cabrini, M., Matti, V., Oldani, A. & di Fagagna, F. D. DICER, DROSHA and DNA damage response RNAs are necessary for the secondary recruitment of DNA damage response factors. *Journal of Cell Science* **129**, 1468-1476, doi:10.1242/jcs.182188 (2016).

13 Eren, M. K., Kilincli, A. & Eren, O. Resveratrol Induced Premature Senescence Is Associated with DNA Damage Mediated SIRT1 and SIRT2 Down-Regulation. *Plos One* **10**, doi:10.1371/journal.pone.0124837 (2015).
